# Cardiomyocyte-expressed TGFβ signals to fibroblasts to program early heart maturation and adult myocyte identity

**DOI:** 10.1101/2025.09.22.677845

**Authors:** Rachel A. Minerath, Rajesh K. Kasam, Casey O. Swoboda, Vikram Prasad, Kelly M. Grimes, N. Scott Blair, Hadi Khalil, Christina M Alfieri, Logan Eads, Anthony J. Saviola, Mohamad Azhar, Lianjie Miao, Mingfu Wu, Michelle Tallquist, Kirk C. Hansen, Matthew T Weirauch, Katherine E. Yutzey, Douglas P. Millay, Jeffery D. Molkentin

## Abstract

Transforming growth factor β (TGFβ) is a secreted growth factor that is sequestered to the extracellular matrix (ECM) as a latent complex. In adult disease TGFβ release in the heart transforms fibroblasts into a differentiated state that synthesizes more ECM. However, it is not known how TGFβ functions in the early developing heart to impact resident fibroblasts. Here, we observe that deletion of the *Tgfb1*, *Tgfb2*, and *Tgfb3* genes (TGFβ ligands) from cardiomyocytes in the early developing heart results in cardiac dysfunction by 6 weeks of age with altered fibroblast activity and altered ECM content. Early postnatal hearts from *Tgfb1/2/3* cardiomyocyte-deleted mice are dysmorphic and cardiac fibroblasts have incorrect activity and produce inappropriate ECM with reduced stiffness. Gene expression profiling of hearts from myocyte-specific *Tgfb1/2/3* deleted mice reveal defects in both cardiomyocyte and fibroblast maturation with ectopic expression of multiple skeletal muscle-specific genes beginning at embryonic day 17.5 and progressing with age. However, cardiomyocyte-specific deletion of TGFβ receptors I/II encoding genes (*Tgfbr1/2*) or Smad2/3 encoding genes (*Smad2/3*) do not recapitulate this phenotype suggesting that TGFβ directly programs early heart fibroblast development that in turn specifies cardiomyocyte maturation. Importantly, *Col1a2^-/-^;Col6a2^-/-^* mice with defective cardiac ECM stiffness, mice lacking cardiomyocyte *Itgb1* with reduced ECM load sensing, and *Tcf21^-/-^* embryos at E17.5 lacking cardiac fibroblasts each fail to generate the same pathologic ECM program with ectopic cardiomyocyte differentiation observed with *Tgfb1/2/3* myocyte-specific deletion. These and additional results indicate that TGFβ generated by cardiomyocytes in the embryonic heart mediates fibroblast differentiation that co-evolves the ECM environment that in turn programs cardiomyocyte maturation to establish their identity.

## INTRODUCTION

Transforming growth factor β (TGFβ) is a secreted growth factor that is sequestered within the extracellular matrix (ECM) as a latent complex in association with latent TGFβ binding protein (LTBP), fibrillin, and often in complex with select integrins on the surface of cells [1]. Activation and release of TGFβ ligand allows it to interact with a heteromeric complex consisting of TGFβ type II receptor (TGFβR2) and TGFβ type I receptor (TGFβR1), inducing downstream signaling through canonical Smad2/Smad3, or through non-canonical signaling [2]. In the diseased adult heart, TGFβ signaling transforms quiescent fibroblasts into myofibroblasts that support cardiac repair and fibrosis through production of ECM and its direct remodeling. Cardiac fibroblast-specific deletion of *Tgfbr1* and *Tgfbr2* or *Smad2* and *Smad3* in the adult heart prevents TGFβ mediated differentiation towards the myofibroblast phenotype, reducing the fibrotic response and cardiac hypertrophy to disease stimuli [3].

Mammals have 3 separate genes encoding TGFβ ligands; *Tgfb1*, *Tgfb2*, and *Tgfb3*. All three genes are expressed in the embryonic heart and regulate crucial developmental processes including epithelial to mesenchymal transition (EMT) and epicardial derived cell (EPDC) invasion of the myocardium, which is the predominant source of fibroblasts in the heart [4–12]. Early studies in *Tgfb1*, *Tgfb2* and *Tgfb3* germline deleted mice showed that all 3 genes are required for heart development or maturation, with loss of the *Tgfb1* gene having the greatest impact on early postnatal cardiomyocytes, which showed greater proliferation and defective ventricular wall maturation [13]. *Tgfb2^-/-^* mice present with malformations of the outflow tract, valve defects, aortic abnormalities, and defective myocardialization [14, 15], while *Tgfb3^-/-^* mice show developmental outflow tract and septal defects, with thickening of heart valves [16].

Cardiomyocyte-localized expression of TGFβ ligands has been observed in the embryonic and injured adult heart suggesting that the cardiomyocyte itself could communicate with cardiac tissue resident fibroblasts [6, 7, 17, 18]. Cardiac fibroblasts develop in concert with cardiomyocytes within the early embryonic and fetal heart. Epicardial derived cells that are transcription factor 21 positive (Tcf21+) and Wilms’ Tumor gene 1 positive (WT1+) migrate into the compact myocardium to generate interspersed tissue resident fibroblasts beginning at embryonic (E) day 12.5 [19, 20]. These newly formed cardiac fibroblasts produce fibronectin, collagen, diverse proteoglycans and heparin-binding EGF-like growth factor as the embryonic heart matures, supporting cardiomyocyte proliferation and subsequent ventricular wall compaction [19, 20]. This maturation of cardiomyocytes likely depends on both paracrine signals and maturation of the ECM environment, given that cardiomyocyte-specific deletion of the gene encoding beta1-integrin in the early mouse embryonic heart results in reduced myocardial cell proliferation and impaired ventricular compaction, suggesting a requirement for force transduction through the ECM for myocyte development [19].

In models of heart development using either cultured embryonic stem (ES) cells or induced pluripotent stem cells (IPSc) to generate engineered heart tissues (EHTs) or embryoid like structures, inclusion of fibroblasts is known to be critical, where they help generate more refined 2D or 3D structures, promote cardiomyocyte maturation with differentiated gene expression, and robustly promote contractile activity [21–24]. Moreover, these *ex vivo* generated artificial cardiac tissues generate peak force when the fibroblasts to cardiomyocytes ratio is 1:1 ratio that approximates the density of these cells in the adult heart [24]. Analysis of engineered heart constructs also suggests that embryonic heart fibroblasts are better than adult heart fibroblasts in promoting cardiomyocyte maturation and augmenting function [25]. Moreover, single-cell RNA sequencing of hearts throughout postnatal development into adulthood identified fibroblast subtype switching and maturation as a major driver of cardiomyocyte maturation [26]. This fibroblast maturation affects ECM content and stiffness that are known to evolve in the early developing heart [27–30]. For example, fibronectin and fibrillin-2 in the embryonic hearts are replaced by fibrillar collagens and laminins in the postnatal period [27–29, 31]. These changes in the postnatal heart are accompanied by increased ECM stiffness that is known to drive cellular differentiation and maturation [29, 30, 32, 33].

Here, we observed that TGFβ is expressed in the embryonic and postnatal heart and downregulated by postnatal (P) day 21. To assess the role of TGFβ during developmental heart maturation the *Tgfb1*, *Tgfb2*, *Tgfb3* genes were deleted in cardiomyocytes of the mouse heart using the α-myosin heavy chain (MHC) promoter. The results indicate that TGFβ generated from the cardiomyocyte compartment in the embryonic hearts regulates resident cardiac fibroblast differentiation and the proper maturation of the ECM, which when deficient results in misdifferentiation of cardiomyocytes and dilated cardiomyopathy and lethality by 5-7 months of age.

## MATERIALS AND METHODS

### Mouse models

*Tgfbr1/2*-loxP targeted mice were previously described [3, 34]. *Smad2/3-* loxP targeted mice were previously described as well [3, 35]. *Tgfb1*-loxP targeted mice [36], *Tgfb2*-loxP targeted mice [37], *Tgfb3*-loxP targeted mice [38] were previously generated. *αMHC-Cre* mice and *αMHC-MerCreMe*r (MCM) mice were used to generate cardiomyocyte specific deletions as previously described [39, 40]. *Tcf21^-/-^* mice and littermate controls for E17.5 RNA sequencing were previously described [20]. *Col1a2^-/-^* mice [41] and *Col6a2^-/-^* mice [42] were previously generated and crossed to generate *Col1a2^-/-^;Col6a2^-/-^* mice. *Itgb1^fl/fl^* mice [43, 44], were crossed with *αMHC-Cre* to generate *Itgb1^fl/fl-αMHC Cre^* mice. Mice were genotyped using the following primers in 5’ to 3’ orientation: *Tgfb1^fl/fl^* (Fwd: aggacctgggttggaagtg, Rev: cttctccgtttctctgtcaccctat), *Tgfb2^fl/fl^* (Fwd: gaaagaaagaaaggacggacctg, Rev: ctgaatgcaagcagttaacccagt), *Tgfb3^fl/fl^* (Fwd: agataaacaatggagtctgtcatgg, Rev1: gtctcatatgtgtcttcctgtctcc, Rev2: ttctggattcatcgactgtgg), *Smad2^fl/fl^* (Fwd: ataatgggtcccagtaacaccggac, Rev: ctcaaaccttaccaagccaaagcag), *Smad3^fl/fl^* (Fwd: gtttactggactgaggttggctgtggc, Rev: gtacagattggccactccggtcactg), *Tgfbr1^fl/fl^* (Fwd: atgagttattagaagttgttt, Rev: accctctcactcttcctgagt), *Tgfbr2^fl/fl^* (Fwd: ctgtgtttactgttaccacaagaaagcagat, Rev: ccagcaaacagagaagagttaactc), *αMHC-Cre* (Fwd: atgacagacagatccctcctatctcc, Rev: ctcatcactcgttgcatcatcgac), *αMHC-MerCreMer* (Fwd: cagggaagtggtggtgtagg, Rev: caggtcgacgatcgtgttggggaagccc), *Col1a2^-/-^* (Fwd: tgtcctgcagaagaagtaaatgaaca, Rev: tgtggtagagggtacattcccatt), Col6a2^-/-^ (Fwd: agctgtctgtttgaatccagccta, Rev WT: caatcaggagcacagaaagaggac, Rev MUT: gtctgtcctagcttcctcactg).

### Tamoxifen treatment of mice

To induce Cre-mediated recombination of loxP containing alleles, mice and controls were injected by intraperitoneal (i.p.) route with pharmaceutical grade tamoxifen (Millipore Sigma, T5648, St. Louis, MO, USA) dissolved in corn oil (80 mg/kg body weight/day). Tamoxifen was solubilized at 37° C with constant agitation for 20 minutes. Injections were administered every other day for a total of 3 injections over 5 days beginning at 4 weeks of age.

### Recombinant TGFβ (rTGFβ) treatment of mice

Recombinant TGFβ (Qkine, Cambridge, UK; Qk010-0050) was reconstituted in 10 mM HCl (Qkine, Qk1001) and administered by intraperitoneal (i.p.) injection daily from postnatal day 1 (P1) to P7 at a dose of 0.5 µg/mouse from P0-P4 and escalated to 1 µg/mouse from P5-P7. Vehicle control mice received equal volumes of 10 mM HCl.

### MyoAAV-A1 production and injection in neonatal mice

To generate MyoAAV-*Tgfb1*, full length *Tgfb1* cDNA from mouse (Dharmacon, MMM1013-202762433) was amplified using iProof High Fidelity DNA polymerase (Bio-Rad, 172-5302) with added restriction overhangs for *AsiSI* and *NheI*. Amplified *Tgfb1* cDNA was ligated into the capsid containing MyoAAV expression vector with a cardiac Troponin T promoter using InFusion® HD Cloning kit. MyoAAV-*Tgfb1* was produced in-house using the MyoAAV 1A capsid as previously described [45–47]. Briefly, recombinant virus was harvested from AAV Pro 293T cells (Takara, Tokyo, Japan) and purified by ultracentrifuge using an iodixanol gradient [47]. AAV titers were quantified using qPCR (Fwd: 5’ gcagtctgggctttcacatg 3’, Rev: 5’ gtgtggcctcagagttttgct 3’). Control virus used in experiments was MyoAAV-luciferase. For early developmental infection, mice were injected at postnatal day 7 (P7) with MyoAAV-Tgfb1 or MyoAAV-luciferase into the thoracic cavity at a dose of 1൷10^12^ genomic copies/mouse.

### Echocardiography

Echocardiography was performed to assess ventricular function and left ventricular dimensions using a Vevo 3100 imaging system with a 40 MHz transducer (VisualSonics, Toronto, ON, Canada). Fur was removed from mice over the imaged area and anesthesia was given to effect with 2% isoflurane inhalation, and mice were subjected to echocardiography in 2D motion (M)-mode. Assessment of heart rate, wall thickness, fractional shortening, and end-diastolic and systolic dimensions were analyzed using the left ventricular (LV) Trace platform in the VevoLab VisualSonics analysis package. Mice were assessed from at least 3 cardiac cycles. Echocardiography and analysis were performed in a blinded manner.

### Ischemia/Reperfusion (I/R) injury

Ischemia/reperfusion injury was performed as previously described [48, 49]. Briefly, 8-week-old mice were anesthetized with 2% isoflurane inhalation to effect and intubated. I/R injury was performed by placing a slipknot around the left coronary artery. After 120 minutes of ischemia, the slipknot was released allowing reperfusion until sacrifice one week later. Mice were treated with sustained-release buprenorphine (0.2 mg/kg) injected subcutaneously for pain management.

### Apical resection model

Apical resection was performed as previously described [50]. Briefly, P7 pups were anesthetized on ice until unresponsive to hind foot pinch. Mice were maintained on ice throughout the procedure to ensure proper anesthesia. The chests of mice were disinfected with 70% ethanol prior to incision. Following skin incision, a lateral thoracotomy was performed, and gentle pressure was applied to exteriorize the heart from the chest cavity. The ventricular apex was then resected up to ∼15% and returned to the chest cavity. Ribs and muscles were sutured to close the chest cavity and skin was closed by applying Gluture topical adhesive (Fisher Scientific, NC0632797, Hampton, NH). Mice were sacrificed at P28 and assessed for regeneration by Masson’s Trichrome staining of heart histological sections.

### RNA extraction and qPCR

Whole ventricles were digested in Trizol (TermoFisher Scientific, Waltham, MA, USA; Cat. #15596018) or cells were scraped in Trizol for mRNA isolation. Trizol-chloroform RNA extraction was performed. Equal amounts of RNA were reverse transcribed to cDNA using iScript cDNA synthesis kit according to manufacturer’s instructions (Bio-Rad, Hercules, CA, USA #1725034). Quantitative real-time PCR was performed using Sso Advanced SYBR green (Bio-Rad, #1725274) on a CFX96 Touch Real-Time PCR System (Bio-Rad). ddCT was used to quantify fold change of target genes normalized to 18S ribosomal RNA expression. Primers used for qPCR analysis are as follows:

### RNA extraction

Whole ventricles were digested in Trizol (TermoFisher Scientific, Waltham, MA, USA; Cat. #15596018) or cells were scraped in Trizol for mRNA isolation. Trizol-chloroform RNA extraction was performed. Equal amounts of RNA were reverse transcribed to cDNA using iScript cDNA synthesis kit according to manufacturer’s instructions (Bio-Rad, Hercules, CA, USA #1725034). Quantitative real-time PCR was performed using Sso Advanced SYBR green (Bio-Rad, #1725274) on a CFX96 Touch Real-Time PCR System (Bio-Rad). ddCT was used to quantify fold change of target genes normalized to 18S ribosomal RNA expression. Primers used for qPCR analysis are as follows in 5’ to 3’ orientation: *Tgfb1* (Fwd: ctttaggaaggacctgggttg Rev: gtgtgtccaggctccaaata), *Tgfb2* (Fwd: atcgtccgctttgatgtctc Rev: tacagttcaatccgctgctc), *Tgfb3* (Fwd: tggaggagaactgctgtgta Rev: cctgagcagaagttggcatag), *18S* (Fwd: tttctcgattccgtgggtgg Rev: tcaatctcgggtggctgaac), Atp2a1 (Fwd: ccatctgcctgtccatgtcc Rev: ggggggtggttatccctccag), *Tnni2* (Fwd: agagtgtgatgctccagatagc Rev: agcaacgtcgatcttcgca), *Mylpf* (Fwd: ttcaaggaggcgttcactgta Rev: tagcgtcgagttcctcattct), *Snai1* (Fwd: cttgtgtctgcacgacctgt Rev: aggagaatggcttctcacca), *Snai2* (Fwd: cattgccttgtgtctgcaag Rev: cagtgagggcaagagaaagg), *Twist* (Fwd: cggacaagctgagcaagat Rev: ggacctggtacaggaagtcg), *Wt1* (Fwd: atccgcaaccaaggatacag Rev: ggtcctcgtgtttgaaggaa), *Acta2* (Fwd: gttcagtggtgcctctgtca Rev: actgggacgacatggaaaag), *Tagln* (Fwd: gactgcacttctcggctcat Rev: ccgaagctactctccttcca), *Cnn1*: (Fwd: gaaggtcaatgagtcaactcagaa Rev: ccatacttggtaatggctttga), *Col1a2* (Fwd: agcaggtccttggaaacctt Rev: aaggagtttcatctggccct), *Col3a1* (Fwd: ctgtaacatggaaactggggaaa Rev: ccatagctgaactgaaaaccacc), *Fn* (Fwd: atgtggacccctcctgatagt Rev: gcccagtgatttcagcaaagg), *Cilp* (Fwd: atggcagcaatcaagacttgg Rev: aggctggactcttctcactga), *Col5a1* (Fwd: cttcgccgctactcctgttc Rev: ccctgagggcaaattgtgaaaa), *Col6a1* (Fwd: ctgctgctacaagcctgct Rev: ccccataaggtttcagcctca), *Ryr1* (Fwd: cagtttttgcggacggatgat, Rev: caccggcctccacagtattg), *Tnni1* (Fwd: atgccggaagttgagaggaaa, Rev: tccgagaggtaacgcacctt), *Tgm2* (Fwd: gacaatgtggaggagggatct, Rev: ctctaggctgagacggtacag), *Fbn2* (Fwd: ctccaccaaagacgctctgg Rev: ccctcgtcccgatactcagg), *Cdkn1a* (Fwd: cctggtgatgtccgacctg Rev: ccatgagcgcatcgcaatc).

### Immunoblotting

For whole tissue protein blots, whole ventricles were excised, washed in cold 1X PBS, and flash frozen. Tissue was homogenized in either 8M Urea in 100 mM ammonium bicarbonate to solubilize ECM proteins or RIPA buffer containing Halt Protease and Phosphatase Inhibitor Cocktail (Thermo Fisher Scientfic, #78442) for total cellular protein blots. For urea protein extract, protein concentration was quantified using a spectrophotometer measuring absorbance at 280 nanometers (A_280_) (Denovix Spectrophotometer/Fluorometer DS-11 FX+, Wilmington, DE). Protein homogenate in RIPA buffer was quantified using a Direct DetectⓇ spectrophotometer (EMD Millipore, Burlington, MA). Laemmli loading buffer was added to samples (0.25% 282 bromophenol blue, 0.5 M dithiothreitol (DTT), 50% glycerol, 10% SDS) and RIPA samples were boiled at 100° C for 5 minutes. For cell lysate, cells were washed in cold 1X PBS and scraped in RIPA buffer containing Halt Protease and Phosphatase Inhibitor Cocktail. Immunoblot samples were prepared as described above. Gel electrophoresis was performed with equal concentrations of protein and transferred to Immobilon-FL PVDF membranes (IPFL00010, EMD Millipore). Nonspecific antibody binding was blocked with 3% (w/v) bovine serum albumin (BSA) diluted in tris-buffered saline (20 mM Tris, 150 mM NaCl, 0.1% (v/v) tween (TBST)) for 1 hour at room temperature. Primary antibody was diluted in blocking buffer, added to the blot, and incubated overnight at 4° C on a rocker. Following incubation in primary antibody, blots were washed 3X with TBST. LI-COR secondary antibody diluted 1:5000 in blocking buffer was added to the blots and incubated for 90 minutes at room temperature on a rocker. Blots were washed 3X in TBST and imaged on a LI-COR Odyssey CLx (LI-COR Biosciences, Lincoln, NE).

### Histology and immunofluorescence staining

Frozen heart sections were used for all histological analysis. Hearts were removed, washed in cold 1X PBS and fixed in 4% v/v paraformaldehyde (PFA) (Electron Microscopy Sciences, Cat#15714, Hatfield, PA) diluted in 1X PBS at 4° C overnight with constant rocking. Embryonic hearts were fixed in 4% v/v PFA for one hour at room temperature. Following fixation, tissues were rinsed with 1X PBS and cryoprotected in 30% sucrose/1X PBS at 4° C overnight. Hearts were embedded in optimal cutting temperature compound (OCT) (Tissue-Tek, #4583, Torrance, CA) and flash frozen in 2-methylbutane pre-chilled on liquid nitrogen. Ten micrometer thick histological sections were generated and washed with 1X PBS to remove OCT prior to staining. Sections were permeabilized and non-specific binding was blocked for 1 hr at room temperature in 2% BSA (w/v) in 0.2% Triton^®^ X-100 (v/v) (Sigma-Aldrich, Cat# 8787, St. Louis, MO) diluted in 1X PBS. Primary antibody was diluted in blocking buffer and incubated overnight at 4° C. The following day, secondary antibody Alexa Fluor 488 or Alexa Fluor 568 secondary antibodies (Invitrogen, Waltham, MA; 1:300) and/or wheat germ agglutinin (WGA) conjugated to Alexa Fluor 488 (ThermoFisher, W11261; 5 ug/mL) were applied at room temperature for 1 hour. Sections were then stained in DAPI (D3571, Invitrogen; Waltham, MA; 1:5000) for 20 minutes at room temperature, washed three times in 1X PBS, and mounted in ProLong Diamond Antifade (P36961, Thermo Fisher Scientific). Immunofluorescent images were acquired using an inverted Nikon A1R confocal microscope and quantified with NIS Elements AR 4.13 software. For general staining of fibrosis, sections were stained with Picrosirius red using the Tissue-Tek Prisma Plus Automated Slide Stainer in the Heart Institute histology core at Cincinnati Children’s Hospital Medical Center or stained for Masson’s trichrome. Images of Picrosirius red and Masson’s trichrome staining were captured using a Leica M165FC stereo microscope with a Leica DFC310 FX camera and the Leica Application Suite. Quantification of cell surface area (CSA) was performed from WGA-stained heart histological sections with ImageJ analysis software (National Institute of Health) by measuring minimally 250 cells/heart.

### Cardiomyocyte isolation and quantification of nucleation

Whole ventricles were washed in cold 1X PBS and fixed in 1% PFA (v/v) at 4⁰ C overnight. The following day, hearts were washed three times in 1X PBS, cut into ∼6 equivalent pieces/heart, and placed into a 2 mL Eppendorf tube containing 2 mg/mL Collagenase B (Roche, #11088807001) diluted in 1X PBS. Hearts were then incubated at 37⁰ C on a shaker. Every 12 hours, collagenase containing isolated cells was collected, added to an equal volume (1.5 mL) of bovine growth serum (Fisher Scientific, # SH3054103), and stored at 4⁰ C. Fresh Collagenase B was added to the remaining heart pieces and incubated as described above. This process was repeated until the heart was digested. For each heart sample, isolated cells were combined for staining. To stain, cells were pelleted by swinging bucket centrifuge at 150x *g* at 4⁰ C. Supernatant was removed and pelleted cells were resuspended in 1X PBS and incubated at room temperature for 5 minutes on a tube rotator. This was repeated two additional times. Following the final wash and centrifugation, non-specific antibody binding was blocked with 2% BSA (w/v) in 0.2% Triton® X-100 (v/v) in PBS and incubated at room temperature for 1 hour on a tube rotator. Cells were pelleted and resuspended in primary antibody for sarcomeric α-actinin (Sigma Aldrich # A7811, 1:100) diluted in blocking buffer and incubated at 4⁰ C overnight on a tube rotator. The next day, cells were washed three times in 1X PBS as described above. Secondary antibody conjugated to Alexa Fluor 488 was diluted 1:200 in blocking buffer, added to pelleted cells, and incubated for 1 hour on a covered tube rotator. Cells were washed one time in 1X PBS and pelleted cells were resuspended in DAPI (diluted 1:5000 in PBS). Cells were incubated at room temperature for 20 minutes on a covered tube rotator. Cells were washed an additional two times in 1X PBS and pellets were resuspended in ProLong Diamond Antifade (P36961, Thermo Fisher Scientific), added to glass slides at a density to achieve isolated cardiomyocytes and covered by coverslip. Immunofluorescent images were acquired by confocal imaging as described above. Nucleation of sarcomeric α-actinin positive cardiomyocytes was quantified using ImageJ (National Institute of Health).

### Collagen hybridizing peptide (CHP) staining

CHP (3Helix, Bio300) staining was performed as previously described [51]. Briefly, fixed tissue sections were blocked with streptavidin and biotin reagents (Thermo Fisher Scientific, E21390) according to the manufacturer’s instructions with incubation in component A for 30 minutes and component B for 30 minutes both at 37⁰ C in a humidified chamber. Following incubation, slides were rinsed two times with 1X PBS for 10 minutes per wash. Non-specific binding was blocked using 5% goat serum (w/v) (Jackson ImmunoResearch #005-000-121; West Grove, PA), 1% BSA (w/v), 0.2% Triton® X-100 (v/v) (Sigma Aldrich, Cat# 8787) in PBS and incubating for 30 minutes at room temperature. A freshly prepared 15 µM CHP solution was warmed for 5 minutes at 80⁰ C, placed on ice for 15 seconds, applied to histological sections, and incubated overnight at 4⁰ C. Sections were washed in 1X PBS and incubated with an Alexa Fluor-488-conjugated secondary streptavidin-conjugated antibody (Invitrogen, S11223).

### Antibodies

Antibodies against the following proteins were used: periostin (Novus Biologicals, Centennial, CO, USA; NBP1-30042; 1:100 dilution for IF, 1:500 dilution for Western blot); Collagen I (Abcam, Boston, MA, USA; ab21286; 1:100 for IF, 1:500 for Western blot), TGFβ1 (Abcam 215715; 1:500 for Western blot), LAP (TGFβ1) (R&D systems, AF-246-NA, Minneapolis, MN; 1:200 for Western Blot), Gapdh (Fitzgerald, #10R-G109A for Western Blot), phospho-SMAD2 (Ser465/467)/phospho-SMAD3 (Ser423/425) (Cell Signaling Technology, #8828, Danvers, MA; 1:500 for Western blot), phospho-SMAD3 (Ser423/425) (Cell Signaling Technology, #9520; 1:500 for Western blot), SMAD3 (Abcam #ab40854; 1:500 for Western blot), SMAD2 (Cell Signaling Technology, # 5339; 1:500 for Western blot), αSMA (Sigma-Aldrich #A2547; 1:100 for IHC), lumican (Abcam #168348; 1:500 for Western Blot), transglutaminase 2 (TGM2, Abcam, #2386; 1:500 for Western Blot), pHH3(Cell Signaling, 9701; 1:100), desmin (Abcam, #15200; 1:100 for IHC), connexin 43 (Cx43, Cell Signaling, 3512; 1:100 for IF), laminin (Sigma-Aldrich, #LO663; 1:100 for IF), SERCA1 (Thermo Scientific, MA3-912, 1:100 for IF), fibronectin (FN, Abcam, #2413; 1:500 for Western Blot), fibrillin-2 (Santa Cruz Biotechnology, Santa Cruz, CA; sc-393968), loxl2 (Invitrogen, PA5-85210), pleiotrophin (Thermo Scientific, PA5-94984), tenascin C (Abcam, #6346), WT1 (Abcam, Ab89901), Cre (EMD Millipore, 69050-3).

### Affymetrix microarray and bioinformatic analysis from adult hearts

Microarray analysis was performed as previously described [52]. Briefly, RNA was extracted from whole hearts of mice at 6 weeks of age. Microarray analysis was performed by the Gene Expression Core Facility (CCHMC) using the Affymetrix Cariom S platform. Differential gene expression was determined by bioinformatic analysis (CEL files) using Transcriptome Analysis Console (Applied Biosystems; ver. 4.0.0.25), Clariom_S_Mouse TAC configuration file (ver. 1), and the iPathway Guide (Advaita Bioinformatics). The raw RNA microarray expression data were submitted to the GEO database and are under review. Accession numbers in the GEO Omnibus database will be provided upon publication acceptance.

### RNA sequencing

RNA sequencing was performed using RNA extracted from isolated fibroblasts from 6-week-old hearts or from whole hearts of mice at embryonic day 17.5 (E17.5) or from whole hearts from mice at postnatal day 7 (P7). RNA isolated from hearts of E17.5 mice and P7 mice was shipped to Novogene (California) and analyzed as previously described [53]. Briefly, paired-end 150 base-pair libraries were generated with NovaSeq 6000 system, and samples were sequenced at a read depth of 40 million reads per sample. Sequencing reads were pseudo-aligned and quantified based using the index transcriptome version of the GRCm39 mouse genome build (NCBI RefSeq GCF_000001635.27). Differentially expressed genes were defined by a p-value <0.05 and a fold change >1.5 or <-1.5. Heatmaps were generated using Morpheus (Morpheus, https://software.broadinstitute.org/morpheus) and value shown are normalized counts. Gene Ontology analysis was performed using the ToppGene Suite [54]. RNA sequencing data of fibroblasts isolated from hearts of 6-week-old *Tgfb1/2/3^fl/f-αMHC-Cre^* mice were processed as previously described [51]. RNA was extracted from fibroblasts using the miRNeasy Micro Kit (Qiagen #217084, Hilden, Germany). RNA sequencing used an Illumina HiSeq2500 instrument within the Human Genetics Department at Cincinnati Children’s Hospital. Differential gene expression cutoffs described above were used. PANTHER analysis was used for gene ontology analysis [55]. The raw RNA sequencing data were submitted to the GEO database and are under review and accession numbers will be provided upon publication acceptance.

### Generation and analysis of single nuclear RNA sequencing

Hearts at postnatal day 7 (P7) were flash-frozen in liquid nitrogen and stored until processing at -80°C. Frozen tissue was pulverized in liquid nitrogen using a pre-cooled mortar and pestle to generate a rough powder, which was then transferred to a sterile 50 mL conical tube maintained on ice. The tissue powder was resuspended in 3mL of pre-chilled Nuclei EZ lysis buffer (Sigma, Product no. N3408 as part of Cat. no. NUC-101) containing 1U/mL of RNAse inhibitor + 0.5% BSA. Sample was dispersed into buffer using a wide-bore 1 mL pipette tip and incubated on ice. After 3 minutes of incubation, 3 mL of 2% BSA in 1X PBS (+1U/mL of RNAse inhibitor) was added to the suspension, and contents gently mixed using a wide-bore 1mL pipette. Contents were transferred to a pre-chilled Dounce homogenizer, and sample homogenized using Pestle A (20-22-strokes). Homogenate/nuclear suspension was then transferred to a pre-chilled 15 mL conical tube and incubated on ice for 10-minutes and subsequently, sequentially filtered through 100, 40, and 20 µm cell strainers and final flow through centrifuged at 500 *xg* for 6 minutes at 4°C. Supernatant was aspirated and the pellet gently resuspended in 2.0 mL of 2% BSA in 1X PBS (+ 0.5U/mL of RNAse inhibitor) and incubated on ice for 5 mins. Sample was then centrifuged at 500 *xg* for 5-minutes at 4°C and supernatant discarded. The nuclear pellet was resuspended in 2 mL of 1% BSA in 1X PBS + 0.5U/mL of RNAse inhibitor + 1:1000 of DAPI [5mg/mL] and incubated for on ice for 10 mins. This nuclear suspension was centrifuged at 500 *xg* for 5 minutes at 4°C and supernatant discarded. Final pellet was resuspended in 0.5 mL of 1% BSA in 1X PBS + 1U/mL of RNAse inhibitor and subjected to flow cytometry sorting for collection of DAPI +ve nuclei. After sorting, the nuclear suspension was made up to 1 mL using 1% BSA in 1X PBS + 1U/mL of RNAse inhibitor, and filtered through a 20 µm cell strainer. Flow through was centrifuged at 1000 *xg* for 5 minutes at 4°C and supernatant discarded. Final nuclear pellet was resuspended in 1% BSA in 1X PBS + 1U/mL of RNAse inhibitor at a concentration of 500-1000 nuclei/mL and sample submitted for visual inspection of nuclear integrity and subsequent 10X nuclear capture. The10X Chromium platform was used to load the nuclei according to the Single Cell 3′ Reagent Kit v3.1, following the guidelines provided by the manufacturer. Around 12,000 nuclei were loaded for each operation. Sequencing was performed on an Illumina NovaSeq 6000 System at the Cincinnati Children’s Hospital Medical Center, department of Human Genetics.

### SnRNA Seq dataset processing

Initial read alignment and quantification of FASTQ files were generated using CellRanger/v5.0.1. For each dataset, we corrected for ambient background RNA using CellBender [56]. Reads from CellBender were then imported into Seurat objects (Seurat/v5.1.0) [57], and nuclei with less than 200 unique features or greater than 20% reads mapping to the mitochondrial genome were removed from downstream analysis. Additionally, only features expressed in at least 3 nuclei from each dataset were retained for downstream analysis. Seurat objects with the remaining nuclei then underwent doublet identification using Solo/v0.2 [58]. Nuclei having higher than the 98th percentile of number of unique features, number of UMIs, and mitochondrial reads after an initial scoring were excluded from downstream analysis. For the purposes of clustering and dimensionality reduction, all datasets were merged and gene count normalization, variable feature identification, and data scaling were performed using the SCTransform function. A PCA was then performed on selected variable features utilizing the RunPCA function, with the npc parameter modified to 60. All datasets were then integrated using the IntegrateLayers function, specifying the normalization method parameter as “SCT” and the method parameter as “CCAIntegration”. and principal component analysis on the variable features was performed using the RunPCA function. To generate the integrated UMAP, dimensionality reduction was performed using 60 principal components using the RunUMAP and FindNeighbors functions. Clustering was performed using the FindClusters function, with a resolution of 0.5. Nuclei predicted to be doublets were then subset and then an initial round of dimensionality reduction with the same parameters was performed. Clusters were then annotated based on known marker genes. If a cluster identified in the first round of dimensionality reduction contained more than 50% of nuclei predicted to be doublets, the cluster was subset prior to rerunning dimensionality reduction functions. The raw sequencing data were submitted to the GEO omnibus and are under review and accession numbers will be provided upon publication.

### Cellchat Analysis

Matrices containing gene counts from cardiomyocyte and FAPs were normalized by the Seurat function NormalizeData and subsequently were used as inputs to create two cell chat objects, representing *Tgfb1/2/3^fl/f^*^l^ and *Tgfb1/2/3^fl/fl-αMHC-Cre^* datasets. The cell chat mouse database was subset to contain only interactions annotated as “ECM-Receptor” interactions. Both cell chat objects were then preprocessed using standard parameters for the functions subset Data, identify OverExpressedGenes, and identify OverExpressedInteractions functions. Inferred communication probabilities for interactions within each genotype were generated using the trimean method in the compute CommunProb function. Cellchat objects were then merged using the merge CellChat function for the purposes of downstream differential interaction strength identification. The compare Interactions function was used on this merged cell chat object to identify the number of interactions and the predicted strength of interactions in the *Tgfb1/2/3^fl/f^*^l^ and *Tgfb1/2/3^fl/fl-αMHC-Cre^* datasets. Differential interactions were identified using the subset Communication function, and interactions identified with stronger representation in *Tgfb1/2/3^fl/f^*^l^ hearts were visualized using the net Visual_bubble functions.

### Isolation and transduction of neonatal cardiomyocytes

Whole hearts were extracted from *Tgfb1/2/3^fl/fl^* hearts at P3. Hearts were briefly rinsed in 1X PBS and atria were removed. Hearts were then transferred to a 50 mL conical Falcon tube containing chilled 1X Ca/Mg-free HBSS (Fisher Scientific; #SH3058801) and kept on ice. Hearts were allowed to settle by gravitational sedimentation and washed twice in Ca/Mg-free HBSS. Hearts were then subjected to trypsin digestion by aspirating off the wash supernatant and adding 9 ml of chilled Ca/Mg-free HBSS. One vial of trypsin (Worthington Biochemical; #LK003225, Lakewood, NJ; ∼560U/vial) was resuspended in 2 ml of chilled Ca/Mg-free HBSS and added to the hearts (11 mL total). Hearts were incubated in trypsin solution at 4° C on nutator for 2 hours with constant, moderate agitation. Following incubation, hearts were allowed to gradually warm at room temperature for 15 minutes. Soybean trypsin inhibitor (Worthington Biochmical; #LK003235) was resuspended in 1 mL of Ca^2+^/Mg^2+^-free HBSS which was then added to trypsin heart mix and incubated at 37⁰ C for 15 minutes on nutator with constant, moderate agitation. Following incubation, one vial of collagenase (Worthington Biochemical; #LK003245, 1600-1800 U/vial) was resuspended in 5 mL pre-warmed Medium L-15 and incubated for 37° C for 1 hour with constant, moderate agitation. Hearts were gently triturated with 10 ml pipette. Hearts were incubated for an additional 15 minutes on nutator at 37° C.

Hearts were triturated until completely dissociated. The volume of cell suspension was made up to 50 mL with M199 media + 15% BGS, mixed gently, and filtered through 70 um sterile cell-filter. Cells were pelleted at 290x *g* at room temperature for 4 minutes. The supernatant was aspirated off and cells were resuspended in M199 media + 15% BGS. Centrifugation was repeated and the cell pellet was resuspended in M199 media + 10%BGS. Centrifugation was performed again and the cell pellet was resuspended in M199 media + 5% BGS. Cells were then plated in 6 well plates pre-coated with 0.1% Gelatin.

The following day, media was replaced with M199+1% BGS. The next day, cells were transduced with previously generated and validated adenovirus (Ad)-GFP (Adenovirus expressing green fluorescent protein (GFP)) or Ad-Cre (Adenovirus expressing Cre recombinase) in M199 + 1% BGS and incubated for 42 hours [3]. Media was replaced with fresh M199 media + 1%BGS and incubated for 48 hours. To harvest, cells media was aspirated off and cells were washed with cold 1X PBS. Cells were then scraped in Trizol for RNA or in RIPA buffer + Halt Protease and Phosphatase Inhibitor Cocktail and processed as described above.

### Flow Cytometry

Whole hearts were isolated from mice at either P7 or 6 weeks of age. Hearts were briefly rinsed in 1X PBS and atria were removed. Whole ventricles were minced on ice using surgical scissors into approximately 2 mm pieces followed by complete mincing of the heart with a razor blade. Each dissociated ventricle was digested in a 12 well tissue culture plate in 1.5 ml DMEM containing 2 mg/mL of Worthington Collagenase type IV (#LS004188), 1.2 U/mL dispase II (Roche, #10165859001, Basel, Switzerland), 0.9 mM CaCl_2_ and 2% fetal bovine serum (FBS) at 37° C for 20 min with gentle rotation followed by manual trituration 15-20 times with a 10 mL serological pipette. Tissues were settled by sedimentation and supernatant was applied to a 40 µm mesh strainer (Fisher Scientific 22-363-547) stored on ice. A volume of 1.5 mL fresh digestion buffer was added to remaining tissue followed by two additional rounds of incubation, trituration, and replacement of remaining tissue with fresh digestion buffer. Subsequent rounds of digestion were triturated with a 5 mL serological pipette for round 2 and a p1000 pipette tip (USA Scientific, Ocala, FL, USA; #1112-1720) through 3 rounds. Pooled supernatants from the 3 rounds of digestion were washed in 1X PBS and centrifuged at 200 x *g* for 20 minutes at 4° C in a swinging bucket rotor centrifuge without brakes. The pellet was resuspended in 1 mL Red Blood Cell Lysis Buffer (Sigma #R7757) for one minute at room temperature. Following incubation, RBC lysis buffer was neutralized by adding 10 mL HBSS and centrifuged at 200 x *g* for 20 minutes at 4° C in swinging bucket rotor centrifuge without brakes. The pellet was resuspended in flow cytometry buffer consisting of 1X HBSS (ThermoFisher Scientific, Waltham, MA, USA; Cat. #14025076) with 2% bovine growth serum and 2 mM EDTA. Purified anti-mouse CD16/32 (1:100) was added to samples to block non-specific binding. Samples were vortexed and incubated at 20°C for 5 minutes. Fluorophore-conjugated antibodies against MEFSK4-allophycocyanin (APC) (Miltenyi Biotec, Waltham, MA, USA; 130-120-802;1:50), CD45-fluorescein isothiocyanate (FITC) (BD Biosciences, Franklin Lakes, NJ, USA; 563890;1:200), and CD31-brilliant violet 421 (BioLegend, San Diego, CA, USA; 102423;1:200) were added to samples and incubated on ice for 30 minutes. Samples were then centrifuged at 200x *g* for 5 minutes and washed with flow cytometry buffer. Centrifugation was repeated and samples were resuspended in flow cytometry buffer and filtered into 5 mL polystyrene round-bottom tubes with a cell strainer cap on ice (Falcon 352236). For flow cytometric quantification of immune cells at P7, CD45-BV510(BioLegend #103138), CD11b-Alexa Fluor 700 (AF700) (BioLegend #101222), Ly6G-BV421(BD Biosciences #562737), Ly6C-peridinin-Chlorophyll-Protein cyanine5.5 (PerCP Cy5.5) (Biolegend #128012), CD64-PE-Cyanine7 (PE-Cy7) (BioLegend #139314), and CCR2-PE (BioLegend #150610) were used. Samples were analyzed using a BD LSR Fortessa 4-laser cytometer running BD FACSDiVa software (BD Biosciences, Franklin Lakes, NJ, USA). Unstained and single color-stained samples were included in all quantitative flow cytometry experiments to perform compensation. Initial gating with forward and side scatter was performed to define single cells as previously described [53]. Cell quantitation was performed using FlowJo software (V10, Tree Star, Inc., Ashland, OR, USA) and total cell number was normalized to tissue mass.

### Atomic force microscopy

Whole mouse hearts were washed in cold 1X PBS. Cross sections were cut from hearts by removing atria and apex. Hearts were fixed in 2% PFA for 2 hours at room temperature. Hearts were washed in 1X PBS three times and cryoprotected in 30% sucrose/1X PBS at 4° C overnight. Hearts were then embedded in OCT as described earlier and flash frozen in 2-methylbutane chilled on liquid nitrogen. Square glass coverslips were coated with 5% gelatin and allowed to dry overnight to allow for better adhesion of the cut histological sections. These heart sections were cut at 20 µm and placed onto 5% gel coated square coverslips which were then placed into 35 mm dishes and allowed to air dry for 20 minutes. Prior to AFM analysis, sections were incubated in 1X PBS for 15 minutes at room temperature. AFM was performed JPK SPM Data Processor Software (Bruker, Billerica, MA) using MLCT Bio Probes, Probe D (Bruker, SKU MLCT-BIO). Force curves were generated in contact mode using the following settings: set point: 1.2, Z-length: 2, Z-speed: 1. The probe was calibrated prior to recording force curves. Analysis of force curves was performed using the JPK Data Processing Program (Bruker).

### ECM mass spectrometry

Preparation of extracellular tissue samples was performed as previously described [59]. Briefly, 5 mg of lyophilized heart tissue was processed in a stepwise extraction with CHAPS and high salt, guanidine hydrochloride, and chemical digestion with hydroxylamine hydrochloride (HA) in Gnd-HCl generating cellular, soluble ECM, and insoluble ECM fractions for each sample. Protein concentration of each fraction was measured using A660 Protein Assay (Pierce, ThermoFisher Scientific, Waltham, MA, USA). Proteolytic digestion using filter-aided sample preparation (FASP) protocol [60] with 10 kDa molecular weight cutoff filters (Sartorius Vivacon 500, Sartorius, Gottingen, Germany; #VN01H02) was performed using 30 µg of protein from each fraction. Samples were reduced using 5 mM tris (2-carboxythyl phosphine), alkylated with 50 mM 2-chloroacetamide, and trypsin digested overnight (enzyme:substrate ratio: 1:100) at 37° C. Peptides were recovered from the filter by successive washes with 0.2% formic acid. Samples containing 10 µg of digested peptides were cleaned using Pierce C18 Spin Tips (ThermoFisher Scientific #84850) according to the manufacturer’s protocol, dried using vacuum centrifugation, and resuspended in 0.1% formic acid diluted in mass spectrometry-grade water.

Liquid chromatography-mass spectrometry (LC-MS/MS) was done using an Easy nLC 1200 instrument coupled to an Orbitrap Fusion Lumos Tribrid mass spectrometer (ThermoFisher Scientific), as previously described [59]. Fragmentation spectra were searched against the UniProt *Mus musculus* proteome database (Proteome ID #UP000000589) using the MSFragger-based FragPipe computational platform [61]. Contaminants and reverse decoys were automatically added to the database. The precursor-ion mass tolerance and fragment-ion mass tolerance were set to 10 ppm and 0.2 kDa, respectively. Fixed modifications were set as carbamidomethyl (C), and oxidation (M), oxidation (P) (hydroxyproline), Gln ->pyro-Glu (N-term), deamidated (NQ), and acetyl (Protein N-term) were set as variable modifications. Two missed tryptic cleavages were permitted. The protein-level false discovery rate (FDR) was ≤1%.

### Di8 Anepps staining

Live cardiomyocytes were isolated from hearts of 8-week-old mice by Langendorff perfusion as previously described [62]. Briefly, mice were injected i.p. with 100 U/mL heparin 10 minutes prior to sacrifice. Hearts were excised, cannulated, and perfused with collagenase digestion buffer. Dissociated hearts were passed through a 100 µm mesh filter and cardiomyocytes were settled for 15 minutes. Sedimented cardiomyocytes were subsequently rinsed, subjected to calcium reintroduction, plated onto laminin coated Ibidi plates (Ibidi #80826-90, Fitchburg, WI) and allowed to attach for 3 hours at 37⁰ C. Following attachment, media was replaced with fresh cell culture media containing 25 µM Di8 Anepps and incubated for 25 minutes at room temperature in the dark on a rocker. Following incubation, cells were washed 5 times in sterile 1X PBS, placed in modified Tyrode’s solution containing 10 mM EGTA, 140 mM LiCl, 4 mM KCl, 1 mM MgCl_2_, 10 mM glucose, 5 mM HEPES (pH 7.4) and immediately imaged by confocal microscopy.

### Transmission Electron Microscopy

Mice were anesthetized and hearts were perfused with 1% paraformaldehyde/2% glutaraldehyde (vol/vol) in cardioplegic solution (50 mmol/L KCl, 5% dextrose in 1X PBS), followed by 1% paraformaldehyde/2% glutaraldehyde (vol/vol) in 0.1 mol/L cacodylate buffer, pH 7.2. Hearts were excised and fixed in 1% paraformaldehyde/2% glutaraldehyde (vol/vol) in 0.1 mol/L cacodylate buffer, pH 7.2 at 4C. Post-fixation was performed in 2% OsO4 (in 0.1 mol/L cacodylate buffer) followed by dehydration in acetone and embedding in epoxy resin. Ultrathin sections were counterstained with uranium and lead salts as described previously [63]. Images were acquired with a Hitachi 7600 electron microscope with an AMT camera.

### Hydroxyproline assays

Hydroxyproline assays were performed as previously described [64]. Briefly, heart tissue was hydrolyzed in 6 N HCl. After neutralization, samples were vacuum dried and resuspended in 5 mM HCl. Chloramine T solution was added to samples and absorbance was assessed at 558 nm.

### Animal welfare and ethics

No human subjects or human tissues were used in this study. Animals were handled in accordance with principles and procedures of the Guide for the Care and Use of Laboratory Animals. All procedures were approved by the Institutional Animal Care and Use Committee (IACUC) at Cincinnati Children’s Hospital Medical Center (Protocol #2021-0047, 2024-0043). Animal groups and experiments were performed in a blinded manner when possible. Age matched littermates and/or Cre controls were used. The number of mice used in this study reflects the minimum number needed to achieve statistical significance based on experience and previous power analysis. Both sexes of mice were used in equal ratios and all animals were housed in a germ-free barrier at 21–22 °C, 40–60% humidity, 12-h light/12-h dark cycle, and unless specified differently below with food and water *ad libitum,* and observed daily by veterinary staff.

### Statistics

Data are expressed as mean +/-SEM. Statistical analysis was performed using GraphPad Prism. For comparison of two groups of normally distributed data, Student’s t-test was performed. For comparison of two groups of non-normally distributed data, Mann Whitney tests were performed. In the case that data was not normally distributed due to large distribution of data within the groups, parametric tests were performed. For comparison of multiple groups, a One-way or Two-way ANOVA was performed with Tukey’s multiple comparisons test. In the case of non-normally distributed data, Kruskal Wallis tests were used with Dunn’s multiple comparisons test. For survival analysis, two-tailed Mantel-Cox tests were performed. Gene input for Gene Ontology enrichment analysis was based on genes that were >1.5 fold upregulated or downregulated with p-value <0.05. Supplementary Figure 2 was subject to Panther analysis and a Fisher Test was used and the correction was FDR <0.05. Supplementary Figure 4b,c was analyzed by ToppFun with a Probability Density Function Test and the correction was FDR <0.05.

## RESULTS

### Analysis of TGFβ expression and deletion of *Tgfb1/2/3* genes in the mouse heart

Previous work suggested that TGFβ expression is preferentially observed during heart development where cardiomyocytes appear to be a significant source [5–7, 17]. We observed prominent TGFβ protein in the embryonic heart at E16.5 days post coital, through postnatal (P) day 7 (P7) (Fig. 1a). However, by P21 TGFβ protein expression was downregulated. To directly examine the functional role of TGFβ ligands in the heart we crossed loxP site targeted alleles for *Tgfb1*(*Tgfb1^fl/fl^*), *Tgfb2* (*Tgfb2^fl/fl^*), and *Tgfb3* (*Tgfb3^fl/fl^*) to generate *Tgfb1/2/3^fl/fl^* mice (Fig. 1b-d) [36–38]. Previous work with germline deletion of *Tgfb1* showed prominent developmental heart effects with altered cardiomyocytes [13]. Isolated neonatal cardiomyocytes from hearts of *Tgfb1/2/3^fl/fl^* mice transduced with adenovirus Cre recombinase (Ad-Cre) displayed significant reductions in *Tgfb1*, *Tgfb2*, and *Tgfb3* mRNA as well as deletion of TGFβ1 by western blot analysis compared to control adenovirus-green fluorescent protein (Ad-GFP) (Fig. 1b,c). The cardiomyocyte-specific αMHC-Cre transgene (TG) was employed, which is expressed as early as embryonic day E8.0 to delete these three genes in vivo (Fig. 1d) [65].

**Fig. 1.**
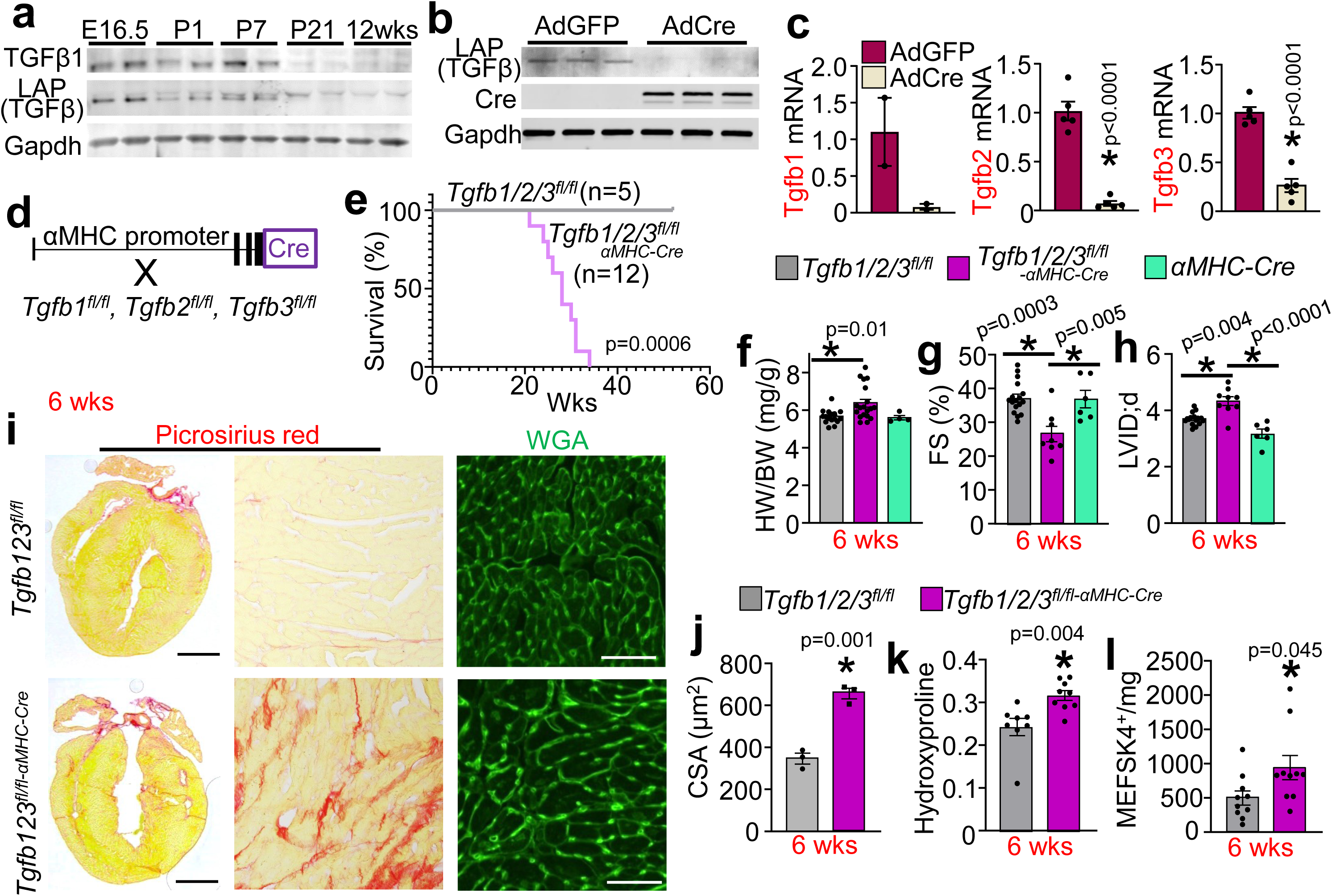
Cardiomyocyte-specific loss of TGFβ ligands results in cardiac dysfunction. **a**. Western blot showing time course of TGFβ1 or Latency associated peptide (LAP) (TGFβ) expression in the heart at embryonic day 16.5 (E16.5), postnatal day 1 (P1), P7, P21, and 12 weeks of age. Gapdh was used to normalize protein loading. **b**. Western blot for TGFβ1 and Cre from protein extracts of neonatal cardiomyocytes from *Tgfb1/2/3^fl/fl-αMHC-Cre^* mice transduced with Ad-GFP or Ad-Cre 48 hrs prior in culture. **c**. mRNA expression of *Tgfb1*, *Tgfb2*, and *Tgfb3* from neonatal cardiomyocytes transduced with Ad-GFP (adenovirus expressing green fluorescent protein) or Ad-Cre (adenovirus expressing Cre recombinase). n=2-5 per treatment group, *p<0.05. **d**. Schematic of the generation of *Tgfb1/2/3^fl/fl-αMHC-Cre^* mice whereby *Tgfb1^fl/fl^*, *Tgfb2^fl/fl^*, and *Tgfb3^fl/fl^* mice were crossed to the *αMHC-Cre* transgene. **e**. Kaplan-Meier curve depicting survival of *Tgfb1/2/3^fl/fl^* and *Tgfb1/2/3^fl/fl-αMHC-Cre^* mice over time in weeks. **f**. Heart-weight to body-weight (HW/BW) ratio of *Tgfb1/2/3^fl/fl^*, *Tgfb1/2/3^fl/fl-αMHC-Cre^*, and *αMHC-Cre* mice at 6 weeks of age. n=4-19 mice in each group, *p<0.05. **g**. Fractional shortening (FS%) at 6 weeks of age of the same groups of mice as in f. n=6-17 mice per group. *p<0.05. **h**. Left ventricular internal diameter in diastole (LVID;d) at 6 weeks of age in the same groups of mice as in f. n=6-17 mice per group. *p<0.05. **i**. Representative histological sections from hearts stained with Sirius Red to assess collagen deposition and wheat germ agglutinin-FITC (WGA, green) to assess cardiomyocyte cross sectional area (CSA) of *Tgfb1/2/3^fl/fl^* and *Tgfb1/2/3^fl/fl-αMHC-Cre^* mice at 6 weeks of age. Scale bar Sirius Red: 2 mm; Scale bar WGA: 100 µm. **j**. Quantification of average cardiomyocyte CSA from WGA staining as shown in i. A minimum of 250 myocytes were measured from each heart, across 3 hearts (points on the graph). *p<0.05. **k**. Hydroxyproline quantification of fibrosis from the left ventricle of hearts at 6 weeks of age as shown in j. n=8-9 mice per group. *p<0.05. **l**. Flow cytometry quantification of MEFSK4+ fibroblasts at 6 weeks of age in the hearts of mice shown in j. n=9-10 mice per group. *p<0.05. c,j,l. Data are presented as mean +/-SEM. Statistical analysis was performed using Student’s T-tests. f-h. Statistical analysis was performed using One Way ANOVAs with Tukey’s multiple comparisons test. k. In the case of non-normally distributed data, Mann Whitney tests were performed. Data are presented as mean +/-SEM.

In previous work, germline deletion of *Tgfb1*, *Tgfb2*, or *Tgfb3* resulted in postnatal mortality between days 15-36 (*Tgfb1^-/-^*) or within hours of birth (*Tgfb2^-/-^* and *Tgfb3^-/-^*) [13, 14, 16]. While *Tgfb1/2/3^fl/fl-αMHC-Cre^* mice are viable, they began to show lethality by 6 months of age compared with two control groups consisting of *Tgfb1/2/3^fl/fl^* and αMHC-Cre TG mice (Fig. 1e). By 6 weeks of age, *Tgfb1/2/3^fl/fl-αMHC-Cre^* mice exhibited a significant increase in heart-weight to body-weight ratio (HW/BW) (Fig. 1f). Echocardiographic analysis revealed a significant reduction in fractional shortening (FS%) with increased left ventricular dilation in diastole (LVID;d) compared with the two control groups (Fig. 1g,h). While TGFβ is an established master regulator of cardiac fibrosis, unexpectedly, *Tgfb1/2/3^fl/fl-αMHC-Cre^* hearts showed increased fibrosis at 6 weeks of age with hypertrophy and increased cardiomyocyte cross-sectional area (Fig. 1i,j). The increase in collagen content was confirmed by hydroxyproline quantification of the left ventricle (Fig. 1k). In addition to fibrosis, *Tgfb1/2/3^fl/fl-αMHC-Cre^* hearts showed expansion of the MEFSK4-positive cardiac fibroblast population at 6 weeks of age as measured by flow cytometry (Fig. 1l).

Hearts of *Tgfb1/2/3^fl/fl-αMHC-Cre^* mice at 6 months of age showed a progressive decline in function and pathology compared to control hearts, such as a significant increase in HW/BW and lung-weight to body-weight ratio (LW/BW) with a significant reduction in FS% and increased LVID;d (Supplementary Fig. 1a-d). As at 6 weeks of age, hearts from 6-month-old *Tgfb1/2/3^fl/fl-αMHC-Cre^* mice showed widespread fibrosis and increased periostin deposition, with progressive cardiomyocyte hypertrophy (Supplementary Fig. 1e). Thus, loss of TGFβ ligands from the cardiomyocyte during early heart development resulted in a progressive decline in cardiac function and fibrosis through adulthood.

### *Tgfb1/2/3* cardiomyocyte deleted mice show altered fibroblasts and fibrosis

Because fibrosis in hearts of *Tgfb1/2/3^fl/fl-αMHC-Cre^* mice was an unexpected finding and TGFβ is a known regulator of fibroblast maturation, we further examined ECM structure-function. Using mass spectrometry protein quantitation from hearts of 6-week-old mice, we observed aberrant upregulation of numerous ECM and matricellular proteins (Fig. 2a). Upregulation of collagen I, lumican, periostin, and transglutaminase 2 were confirmed by western blot analysis (Fig. 2b). Additionally, hearts from *Tgfb1/2/3^fl/fl-αMHC-Cre^* mice exhibited increased collagen I and periostin deposition as confirmed by immunohistochemistry at 6 weeks of age (Fig 2c-e). Given the increase in select ECM proteins we next examined the organization of the ECM in hearts from *Tgfb1/2/3^fl/fl-αMHC-Cre^* mice. Ischemia-reperfusion injury was employed along with histological processing and analysis by transmission electron microscopy (TEM), which showed that the scar area and collagen fibrils from hearts of *Tgfb1/2/3^fl/fl^* control mice was highly aligned and dense but the scar area in hearts from *Tgfb1/2/3^fl/fl-αMHC-Cre^* mice displayed collagen fibers that were shorter, sparse, and less organized (Fig. 2f).

**Fig. 2.**
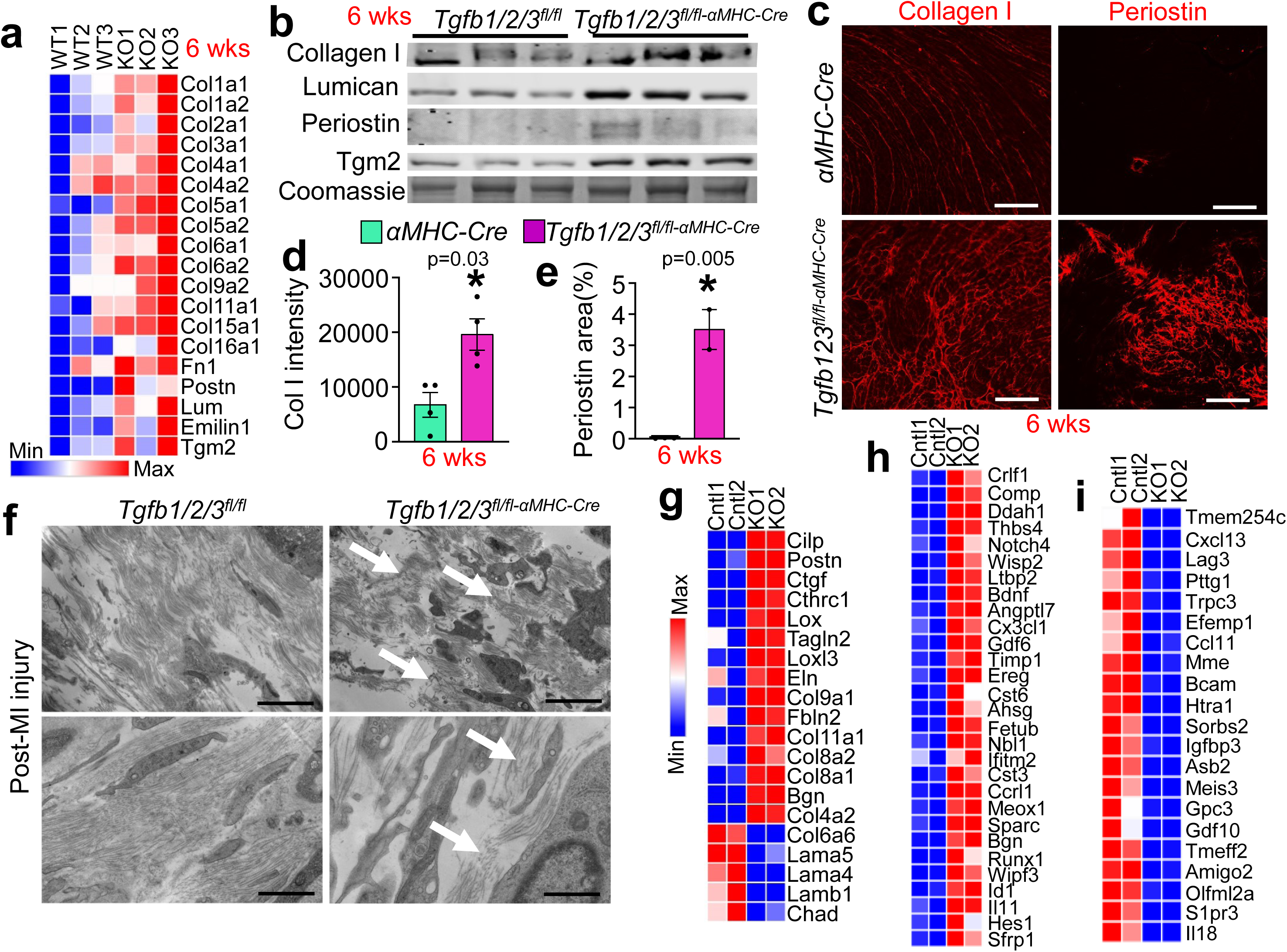
Hearts of *Tgfb1/2/3^fl/fl-αMHC-Cre^* mice exhibit fibrosis, altered extracellular matrix (ECM) and deficient fibroblast maturation. **a**. Heatmap of select dysregulated proteins from cardiac ECM preparations by mass spectrometry performed at 6 weeks of age *Tgfb1/2/3^fl/fl^* and *Tgfb1/2/3^fl/fl-αMHC-Cre^* mice. **b**. Western blot validation of select proteins at 6 weeks of age from hearts of mice shown in a. Coomassie is shown as a loading control. **c**. Representative immunohistochemistry images for collagen I (red, left panel) and periostin (red, right panel) expression from hearts of αMHC-Cre and *Tgfb1/2/3^fl/fl-αMHC-Cre^* mice at 6 weeks of age. Scale bars, periostin: 300 µm. Scale bars, collagen: 200 µm. **d**. Quantification of collagen I staining from images as shown in C. n=4 hearts per group. *p<0.05. **e**. Quantification of periostin staining from images as shown in C. n=2-3 hearts per group. *p<0.05. **f**. Representative images of transmission electron microscopy (TEM) from the scar area of hearts of *Tgfb1/2/3^fl/fl^* and *Tgfb1/2/3^fl/fl-αMHC-Cre^* mice subjected to ischemic injury for 2 hours followed by reperfusion. Mice were sacrificed 1-week post-injury and the scar area was imaged by TEM. Scale Bars: 2 µm. Arrows depict regions of sparse and disorganized collagen. **g**. Heatmap from RNA sequencing for ECM, matricellular and attachment genes performed on cardiac fibroblasts isolated from hearts of *Tgfb1/2/3^fl/fl^* and *Tgfb1/2/3^fl/fl-αMHC-Cre^* mice at 6 weeks of age. **h**. Heatmap from RNA sequencing showing induced genes with unique or unexpected functions in cardiac fibroblasts isolated from hearts of *Tgfb1/2/3^fl/fl^* and *Tgfb1/2/3^fl/fl-αMHC-Cre^* mice at 6 weeks of age. See Supplementary Table 1 for annotation of gene function. **i**. Heatmap from RNA sequencing showing repressed genes with unique or unexpected functions in cardiac fibroblasts isolated from hearts of *Tgfb1/2/3^fl/fl^* and *Tgfb1/2/3^fl/fl-αMHC-Cre^* mice at 6 weeks of age. See Supplementary Table 1 for annotation of gene function. d,e. Data presented as mean +/-SEM. Student’s T-tests were used.

To assess how the loss of TGFβ ligands from cardiomyocytes alters fibroblast differentiation, we performed RNA sequencing of fibroblasts isolated from 6-week-old hearts from *Tgfb1/2/3^fl/fl-αMHC-Cre^* mice versus controls. Differential expression analysis showed a pro-fibrotic and altered signaling profile in fibroblasts from hearts of *Tgfb1/2/3^fl/fl-αMHC-Cre^* mice compared to controls (Fig. 2g). Among the most highly upregulated gene ontology (GO) terms were pathways involved in ECM assembly, negative regulation of adhesion and inflammatory responses (Supplementary Fig. 2a). Interestingly, cardiac fibroblasts from hearts of *Tgfb1/2/3^fl/fl-αMHC-Cre^* mice displayed a downregulation of basement membrane genes involved in cellular adhesion structural support including *Lama5*, *Lama4*, *Lamb1*, and *Col6a6* (Fig. 2g). GO terms downregulated in fibroblasts from hearts of *Tgfb1/2/3^fl/fl-αMHC-Cre^* mice represented categories involved in cellular adhesion and cellular differentiation (Supplementary Fig. 2b).

Furthermore, fibroblasts isolated from hearts of *Tgfb1/2/3^fl/fl-αMHC-Cre^* mice showed both upregulated and downregulated expression of several unique genes including growth factors, chemokines, transcription factors, matricellular proteins, and proteins involved in ECM-cellular crosstalk, adhesion and epithelial mesenchymal transition (Fig. 2h,i). In fact, many of these up and down-regulated gene expression changes were unique to the hearts of *Tgfb1/2/3^fl/fl-αMHC-Cre^* mice and not observed in other mouse models of fibrotic heart disease [66, 67].

Supplementary Table 1 describes the function and unique aspects of 43 gene changes in adult heart fibroblasts from *Tgfb1/2/3^fl/fl-αMHC-Cre^* mice. In addition to being unique, some of the differentially expressed genes are also observed in cancer associated fibroblasts or in embryonic mesenchymal cells [68, 69] (Supplementary Table 1). Thus, without cardiomyocyte generated TGFβ in the early developing heart, resident fibroblasts aberrantly differentiate into an altered phenotypic state that resembles a more nascent mesenchymal cell-type.

### Altered cardiomyocyte maturation in heart-specific *Tgfb1/2/3* deleted mice

This altered phenotypic state of cardiac fibroblasts appears to also negatively impact the differentiation of cardiomyocytes in the heart, which we further assessed in hearts of 6-week-old *Tgfb1/2/3^fl/fl-αMHC-Cre^* mice versus controls by bulk RNA sequencing. Strikingly, these hearts displayed a gene expression profile that not only suggested impaired cardiomyocyte maturation, but it also showed ectopic expression of several skeletal muscle-specific gene isoforms including *Atp2a1*, *Tnni2*, *Mylpf*, *Tnnt3*, *Acta1*, *Ankrd1*, *Casq1*, and *Ryr1* (Fig. 3a). A very similar skeletal muscle ectopic gene program has been observed previously in mice with cardiac-specific deletion of key chromatin and epigenetic regulator factors in the embryonic heart (Supplementary Table 2, [70–73]). RNA expression results were confirmed by qPCR analysis against a panel of skeletal muscle-specific isoforms: *Atp2a1*, *Tnni2*, and *Mylpf* (Figure 3b-d). Interestingly, *Atp2a1* (Serca1) protein expression by immunohistochemistry was observed in the hearts of *Tgfb1/2/3^fl/fl-αMHC-Cre^* mice up to 6 months of age (Fig. 3i).

**Fig. 3.**
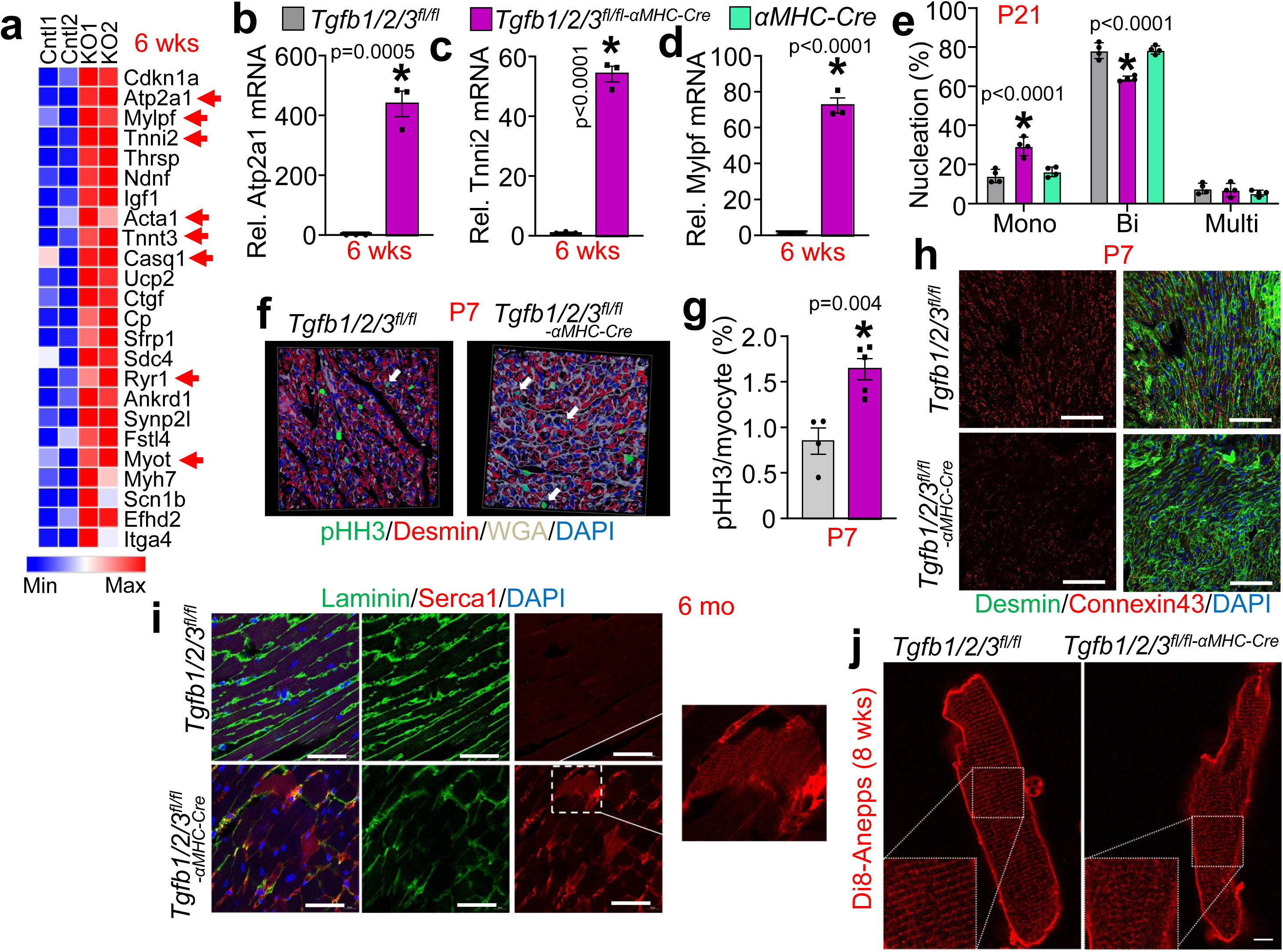
Hearts of *Tgfb1/2/3^fl/fl-αMHC-Cre^* mice at 6 weeks of age show lack of cardiomyocyte maturation and induction of the skeletal muscle gene program. **a**. Heatmap of aberrant gene induction whole heart RNA sequencing performed at 6 weeks of age from *Tgfb1/2/3^fl/fl^* and *Tgfb1/2/3^fl/fl-αMHC-Cre^* mice. The arrows show skeletal muscle specific genes that are ectopically induced. **b**. Quantitative PCR validation of *Atp2a1* at 6 weeks of age from hearts of *Tgfb1/2/3^fl/fl^* and *Tgfb1/2/3^fl/fl-αMHC-Cre^* mice. n=3 mice per group. *p<0.05 **c**. Quantitative PCR validation of *Tnni2* at 6 weeks of age from hearts of *Tgfb1/2/3^fl/fl^* and *Tgfb1/2/3^fl/fl-αMHC-Cre^* mice. n=3 mice per group. *p<0.05. **d**. Quantitative PCR validation of *Mylpf* at 6 weeks of age from hearts of *Tgfb1/2/3^fl/fl^* and *Tgfb1/2/3^fl/fl-αMHC-Cre^* mice. n=3 mice per group. *p<0.05. **e**. Quantification of cardiomyocyte mononucleation, binucleation, and multinucleation from cardiomyocytes isolated from postnatal day 21 (P21) hearts of *Tgfb1/2/3^fl/fl^,* αMHC-Cre, and *Tgfb1/2/3^fl/fl-αMHC-Cre^* mice. Nucleation was counted from 4 hearts in each group with 250 cells per heart. n=4 mice per group. *p<0.05. **f**. Representative image of immunohistochemistry assessment of phospho-histone H3 from the heart at P7 for each indicated genotypes of mice. pHH3-positive cardiomyocytes (white arrows) were assessed by counting pHH3 positive (green) nuclei (blue, 4′,6-diamidino-2-phenylindole (DAPI)). Cardiomyocytes were identified as desmin positive cells (red) outlined by WGA staining (white). **g**. Quantification of phospho-histone H3 positive cardiomyocyte nuclei for each indicated genotype as shown in f. n=4-5 mice per group. *p<0.05. **h**. Representative immunofluorescent images of histological heart sections at P7 for the two genotypes shown for connexin 43 (red) and cardiomyocyte organization using desmin (green). The left panel depicts only connexin 43 staining (red). The right panel merges desmin staining (green) with connexin 43 and DAPI nuclei staining (blue). Scale bar: 40 µm. **i**. Immunohistological analysis of Serca1 expression (red) at 6 months of age from heart histological sections of the two genotypes of mice shown. The middle panel depicts laminin (green) to outline cardiomyocytes but it also shows disorganization in the outer membrane in hearts from *Tgfb1/2/3^fl/fl-αMHC-Cre^* mice. The right panel shows staining of Serca1 (red) being incorporated into the cardiomyocyte in only hearts of *Tgfb1/2/3^fl/fl-αMHC-Cre^* mice, with a 10x increased magnification image. The left panel is a merge of all channels including DAPI. Scale bar: 40 µm. **j**. Confocal imaging of Di8-Anepps staining (red) of live cardiomyocytes in culture at 8 weeks of age from hearts of the indicated genotypes of mice. Scale bar: 5 µm. b-d, g. Data presented as mean +/-SEM. Student’s T-test was used for statistical analysis. e. Data presented as mean +/-SEM. Statistical analysis was performed using Two Way ANOVAs with Tukey’s multiple comparison tests.

Given the observed transcriptional alterations that indicated impaired and ectopically altered cardiomyocyte identity/maturation, we assessed additional maturation markers to understand the timing and extent of the defect. Indeed, cardiomyocytes from hearts of *Tgfb1/2/3^fl/fl-αMHC-Cre^* mice at P21 showed increased mononucleation with reduced binucleation compared to controls (Fig. 3e). Cardiomyocytes from hearts of *Tgfb1/2/3^fl/fl-αMHC-Cre^* mice at P7 also had increased cell cycle activity as measured by phospho-histone H3 (pHH3) (Fig, 3f,g). However, apical resection of the hearts from *Tgfb1/2/3^fl/fl-αMHC-Cre^* mice at P7 was not of sufficient effect to enable regeneration of the heart as assessed at P28 (Supplementary Fig. 3a,b). As another marker of heart maturation, immunohistochemistry for connexin 43 expression in the hearts of *Tgfb1/2/3^fl/fl-αMHC-Cre^* mice at P7 showed reduced signal as well as impaired organization of desmin, which is another marker of maturation (Fig. 3h). Laminin (green) also showed disorganization in the hearts of *Tgfb1/2/3^fl/fl-αMHC-Cre^* mice versus *Tgfb1/2/3^fl/fl^* mice at 6 months of age in histological sections, which we also stained for skeletal muscle specific Serca1 protein to show its prominent ectopic expression in the adult cardiomyocytes (Fig. 3i). Finally, T-tubules are established after birth and continue to mature into adulthood where they eventually display a striated pattern in the mature cardiomyocytes [74]. Histological analysis from hearts of *Tgfb1/2/3^fl/fl-αMHC-Cre^* mice at 8 weeks of age showed reduced T-tubule complexity by di8Anepps staining further indicating impaired cardiomyocyte maturation (Fig. 3j). TEM in hearts of *Tgfb1/2/3^fl/fl-αMHC-Cre^* mice at 5 months of age showed reduced integrity of the intercalated discs and aberrant mitochondrial morphology compared to control hearts (Supplementary Fig. 3c). These findings demonstrate that the loss of TGFβ ligands from the cardiomyocyte results in impaired and improper cardiomyocyte maturation, which is maintained throughout the lifespan of these mice as they progressively succumb to heart failure.

Additional temporal analysis in hearts of *Tgfb1/2/3^fl/fl-αMHC-Cre^* mice at P1 and P7 showed dysmorphic hearts and increased HW/BW (Figure 4a,b). However, at P7 hearts of *Tgfb1/2/3^fl/fl-αMHC-Cre^* mice did not yet display alterations in collagen content nor increased fibroblast numbers (Fig. 4c,d), although atomic force microscopy showed a significant reduction in heart stiffness compared to litter mate controls (Fig. 4e). Analysis of collagen integrity in the hearts of *Tgfb1/2/3^fl/fl-αMHC-Cre^* mice at 4 weeks with collagen hybridizing peptide (CHP) also showed defective ECM maturation compared to control hearts (Supplementary Fig. 3d). Taken together these results suggest that TGFβ is required in the early developing heart to generate a rigorous structural ECM network to support cardiomyocyte differentiation. Indeed, increasing ECM stiffness is known to drive cellular maturation of cardiomyocytes in culture or in engineered heart tissue [29, 30, 32, 33], and engineered heart tissue requires fibroblast co-culture to fully differentiate [21, 23, 24]. In support of these results, hearts from *Tgfb1/2/3^fl/fl-αMHC-Cre^* mice also fail to downregulate embryonic expressed fibronectin and fibrillin-2 (Fig. 4f). Additionally, ECM or matricellular proteins such as Tgm2, Loxl2, pleiotrophin, lumican, and tenascin C in hearts of *Tgfb1/2/3^fl/fl-αMHC-Cre^* mice at P7 were not reduced compared to control hearts (Fig. 4f). Thus, TGFβ generated by cardiomyocyte in the developing heart is required to drive fibroblast and postnatal ECM maturation.

**Fig. 4.**
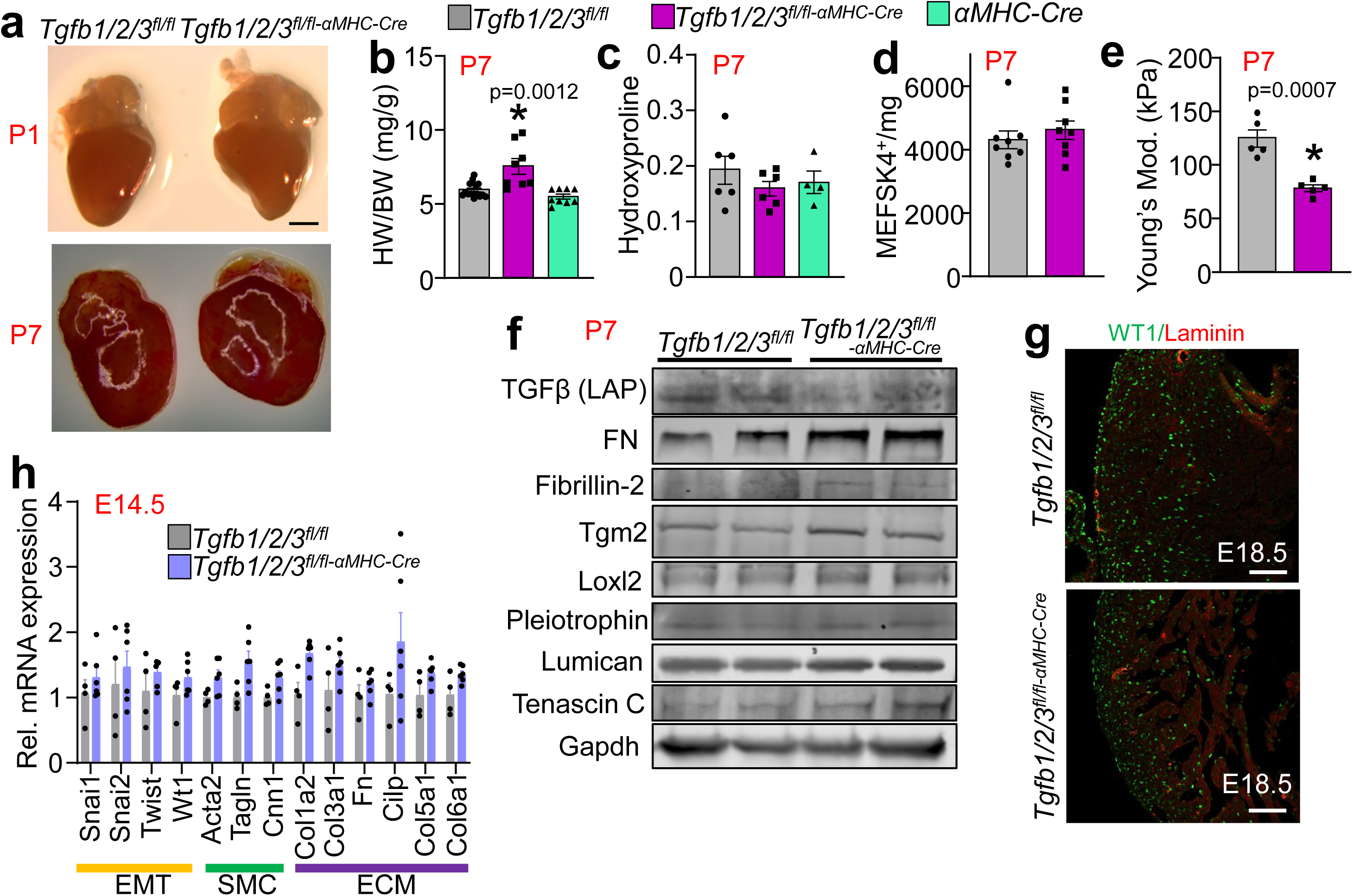
Hearts of *Tgfb1/2/3^fl/fl-αMHC-Cre^* mice are developmentally altered with immature ECM but no changes in epicardial EMT or EPDC migration. **a**. Images of hearts at P1 and P7 from the two genotypes of mice demonstrating early dysmorphia. Scale bar: 1 mm (equal aspect ratio in both pictures). **b**. Heart-weight to body weight ratio (HW/BW) at P7 in the three genotypes of mice shown. n=8-14 mice per group. *p<0.05. **c**. Hydroxyproline quantification of fibrosis from left ventricles of mice at P7 in the three genotypes of mice shown. n=4-6 mice per group. **d**. Flow cytometric quantification of MEFSK4+ fibroblasts from left ventricles of mice at P7 in the two genotypes of mice shown. n=8 mice per group. **e**. Atomic force microscopy quantification of cardiac stiffness at P7 in hearts of *Tgfb1/2/3^fl/fl^* and *Tgfb1/2/3^fl/fl-αMHC-Cre^* mice. A total of 64 force curves were generated from histological sections in minimally 10 different areas of the LV free wall (minimally 640 force curves analyzed/heart). Each datum point depicted represents the average of the force curves generated for a given heart. n=5 mice per group. *p<0.05. **f**. Western blot assessment of extracellular matrix proteins in the hearts of *Tgfb1/2/3^fl/fl^* and *Tgfb1/2/3^fl/fl-αMHC-Cre^* mice at P7. GAPDH was used as a processing and loading control. **g**. Representative immunostaining of cardiac histological sections for WT1 (green) merged with laminin (red) to assess epicardial derived cells migrating into the myocardium at E18.5 for the two genotypes shown. Scale bars: 100 µm. **h**. mRNA expression of genes involved in epithelial-mesenchymal transition (EMT), smooth muscle cell (SMC) maturation, and extracellular matrix (ECM) expression in E14.5 hearts of *Tgfb1/2/3^fl/fl^* and *Tgfb1/2/3^fl/fl-αMHC-Cre^* mice. Genes were normalized to 18S ribosomal RNA expression. n=4-6 mice per group. b,c. Data presented as mean +/-SEM. Kruskal Wallis tests were used with Dunn’s multiple comparisons test for statistical analysis. d,e,h. Data presented as mean +/-SEM. Student’s T-test used for statistical analysis.

We also examined the state of epicardial cells in hearts from *Tgfb1/2/3^fl/fl-αMHC-Cre^* mice, as their migration into the myocardium is critical for heart development and proper fibroblast formation, and TGFβ is an established regulator of both epicardial-mesenchymal transition (EMT) and epicardium-derived progenitor cell (EPDC) migration [8–12]. Using WT1 as a marker of EPDCs, we confirmed by immunofluorescence that hearts from *Tgfb1/2/3^fl/fl-αMHC-Cre^* mice at E18.5 did not display defects in EPDC migration compared to Cre-negative littermates (Fig. 4g). Analysis at an even earlier timepoint by qPCR of hearts from *Tgfb1/2/3^fl/fl-αMHC-Cre^* embryos at E14.5 revealed no changes in expression of genes related to EMT induction, smooth muscle cell (SMC) function, nor ECM function suggesting that EMT and early cell differentiation were unaffected by the embryonic deletion of TGFβ ligands from cardiomyocytes (Fig. 4h). Thus, epicardial EMT and EPDC migration occurs properly, suggesting that the defect observed in cardiac fibroblasts and ECM composition during postnatal heart maturation is due to other factors.

To further examine the potential mechanisms associated with impaired cardiomyocyte maturation, we performed RNA sequencing at multiple embryonic and postnatal time points, beginning at E17.5. Indeed, at E17.5 we observed several differentially regulated genes that are similarly dysregulated in hearts from 6-week-old *Tgfb1/2/3^fl/fl-αMHC-Cre^* mice including *Ryr1*, *Casq1*, *Cdkn1a*, *Sdc4*, *Nppb*, and *Ankrd1* (Supplementary Fig. 4a). Pathway analysis of upregulated genes revealed overrepresentation in ECM production and organization genes (Supplementary Fig. 4b). Downregulated pathways involved cardiac conduction and ion channel activity, heart contraction, and cell junction organization, demonstrating a loss of cardiomyocyte maturation within the embryonic heart (Supplementary Fig. 4c). These included genes such as *Hcn4*, *Hcn1*, *Kcnh7*, *Kcnd3*, *Kcnj3, Tnnt3, Itgad, Mybphl, Myl7,* and *Efemp1*.

However, the profile of ectopic expression of skeletal muscle genes only included *Ryr1* and *Casq1* at E17.5, while by 6 weeks of age many more ectopic genes appeared (Fig. 3a). We also performed RNA-seq analysis in hearts of *Tgfb1/2/3^fl/fl-αMHC-Cre^* mice and controls at P7 and 4 weeks of age (Supplementary Fig. 4d-g). The data show *Ryr1*, *Serca1*, *Mylpf* and *Tnni2* were strongly induced at 4 weeks of age, but only *Ryr1* and *Mylpf* were induced by P7 versus controls. Thus, embryonic hearts from *Tgfb1/2/3^fl/fl-αMHC-Cre^* mice display widespread gene expression alterations that progress to a pathologic state by P7, which intensifies with postnatal heart maturation until early adulthood. These results are reminiscent of developmental epigenetic induction of many of the same skeletal muscle genes in the hearts of mice deleted for chromatin regulatory factors (*Jarid2, Hdac1/2, Chd4, Ezh2)*, which were maintained in adulthood as well (Supplementary Table 2).

### Analysis of TGFβ function during early heart development in *Tgfb1/2/3* deleted mice

We next sought to determine if the maturation effect associated with deletion of TGFβ ligands in the heart was limited to the fetal and early postnatal time windows, or if it continued into adulthood (Supplementary Fig. 5a). *Tgfb1/2/3^fl/fl-αMHC-MCM^* mice or controls were injected i.p. with 80 mg/kg Tamoxifen at 4 weeks of age and analysis was performed 16 weeks later (Supplementary Fig. 5b). Analysis of the *Tgfb1/2/3^fl/fl-αMHC-MCM^* mice showed no changes in cardiac fibrosis, hypertrophy as assessed by HW/BW, or altered structure-function as assessed by echocardiography (Supplementary Fig. 5c-f). There were also no observed changes in fibroblast number measured by flow cytometry quantification of MEFSK4+ cells (Supplementary Fig. 5g). Lastly, hearts from these*Tgfb1/2/3^fl/fl-αMHC-MCM^* adult-deleted mice exhibited no ectopic induction of the skeletal muscle genes *Atp2a1*, *Tnni2*, and *Mylpf*. (Supplementary Fig. 5h-j) These results demonstrate that TGFβ ligands made by cardiomyocytes regulate heart maturation only during early heart development, but not in the adult heart.

While our primary mouse model for analysis resulted in simultaneous deletion of all three *Tgfb* genes, we also assessed the potential for functional redundancy, or if one ligand was more critical than the other two. Hence, we generated TGFβ single gene-deleted mice (*Tgfb1^fl/fl-^ _αMHC-Cre_*_, *Tgfb2*_*_fl/fl-αMHC-Cre_*_, and *Tgfb3*_*_fl/fl-αMHC-Cre_*_) and double deleted mice (*Tgfb2/3*_*_fl/fl-αMHC-Cre_, Tgfb1/3^fl/fl-αMHC-Cre^*, and *Tgfb1/2^fl/fl-αMHC-Cre^*). We assessed cardiac fibrosis of single deleted mice at 6 weeks of age using picrosirius red staining of cardiac histological sections, which showed that only hearts from*Tgfb1^fl/fl-αMHC-Cre^* mice displayed fibrosis similar to hearts from *Tgfb1/2/3^fl/fl-αMHC-Cre^* mice, while hearts from *Tgfb2^fl/fl-αMHC-Cre^* and *Tgfb3^fl/fl-αMHC-Cre^* mice displayed no apparent pathological fibrosis (Fig. 5a). Echocardiographic analysis was performed at 6 weeks of age and only mice lacking *Tgfb1* (*Tgfb1/2/3^fl/fl-αMHC-Cre^*, *Tgfb1^fl/fl-αMHC-Cre^*, *Tgfb1/3^fl/fl-αMHC-Cre^*, and *Tgfb1/2^fl/fl-αMHC-Cre^*) displayed significant reductions in FS%, while mice with genotypes expressing *Tgfb1* did not show cardiac dysfunction (Fig. 5b). Similarly, only mice with combinatorial alleles containing *Tgfb1* deletion displayed ectopic expression of skeletal muscle-specific genes *Atp2a1*, *Mylpf*, and *Tnni2* (Fig. 5c-e). These results indicate that it’s the loss of *Tgfb1* that drives the pathologic phenotype of *Tgfb1/2/3^fl/fl-αMHC-Cre^* mice. However, we were concerned about the potential for intermittent compensation by *Tgfb2* or *Tgfb3*, hence we maintained consistency throughout this study by always using the triple gene deletion strategy.

**Fig. 5.**
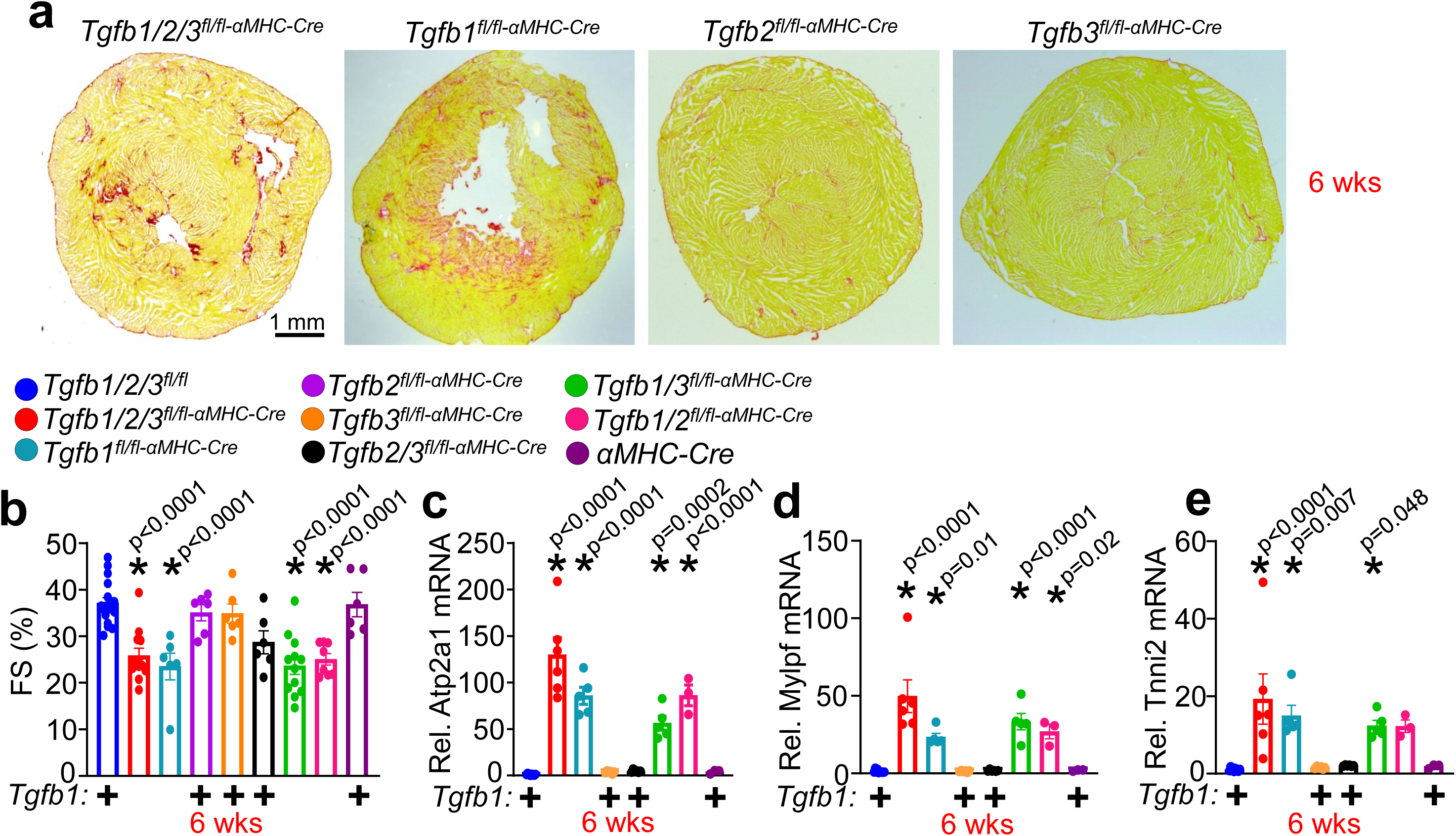
*Tgfb1* drives cardiomyocyte maturation but not *Tgfb2* or *Tgfb3*. **a**. Representative histological images with Sirius red staining (fibrosis) from cross sections of _hearts from *Tgfb1/2/3*_*_fl/fl-αMHC-Cre_*_, *Tgfb1*_*_fl/fl-αMHC-Cre_*_, *Tgfb2*_*_fl/fl-αMHC-Cre_*_, and *Tgfb3*_*_fl/fl-αMHC-Cre_* _mice at_ 6 weeks of age. Scale bar: 1 mm. **b**. Echocardiography measured fractional shortening (FS%) of hearts from mice with the indicated genotypes at 6 weeks of age. The “+” indicates the presence of the *Tgfb1* allele in the different genotypes shown. n=6-17 mice per group. *p<0.05. **c-e**. Quantitative PCR assessment of skeletal muscle-specific genes *Atp2a1*, *Mylpf*, and *Tnni2* from hearts of mice with the indicated genotypes of mice at 6 weeks of age. Relative gene expression is standardized to 18S ribosomal RNA expression. The “+” depicts the presence of the*Tgfb1* allele in the different genotypes shown. n=3-6 mice per group. *p<0.05. b-e. Data are presented as mean +/-SEM. One-way ANOVAs with Tukey’s multiple comparison tests were used to assess statistical significance.

We next examined whether overexpressing *Tgfb1* in the *Tgfb1/2/3^fl/fl-αMHC-Cre^* heart would rescue cardiomyocyte maturation and cardiac dysfunction. To do so, we generated MyoAAV’s to overexpress *Tgfb1* or luciferase under the control of the cardiac troponin T promoter (Supplementary Fig. 6a). Following injection of neonatal mice at P7, mRNA and western blot analysis confirmed upregulation of *Tgfb1* in the heart (Supplementary Fig. 6b,c). Cardiac function was assessed by echocardiography at 6 weeks of age. MyoAAV-Luciferase injected into *Tgfb1/2/3^fl/fl-αMHC-Cre^* mice continue to show a significant reduction in FS% in comparison to MyoAAV-Luciferase injected into *Tgfb1/2/3^fl/fl^* control littermates (Supplementary Fig. 6d).

However, MyoAAV-Tgfb1 injected into *Tgfb1/2/3^fl/fl-αMHC-Cre^* mice conferred no functional benefit over *Tgfb1/2/3^fl/fl-αMHC-Cre^* mice injected with MyoAAV-Luciferase (Supplementary Fig. 6d), nor did it reduce fibrosis assessed by periostin immunostaining or cellular hypertrophy assessed by WGA staining in cardiac histological sections (Supplementary Fig. 6e). Lastly, injection of MyoAAV-Tgfb1 at P7 in *Tgfb1/2/3^fl/fl-αMHC-Cre^* mice did not correct the ectopic induction of *Atp2a1*, *Tnni2*, or *Mylpf* at 6 weeks of age in the heart (Supplementary Fig. 6f-h).

Given that AAV gene transduction requires at least one week to achieve expression in the heart, the current rescue strategy may have been too late in postnatal development to have an effect. Hence, we also performed direct injection of recombinant TGFβ1 (rTGFβ1) at P1 through P7 in pups from *Tgfb1/2/3^fl/fl-αMHC-Cre^* mice versus vehicle control (Supplementary Fig. 7a). Injection of rTGFβ1 was sufficient to induce downstream signaling measured by phospho-Smad2 and phospho-Smad3, as well as increase HW/BW at 6 weeks of age, but it did not improve cardiac function measured by echocardiography (Supplementary Fig. 7b-d). rTGFβ1 treatment did not alter periostin induction in hearts form *Tgfb1/2/3^fl/fl-αMHC-Cre^* mice nor alter the ectopic induction of *Atp2a1*, *Mylpf*, nor *Tnni2* gene expression (Supplementary Fig. 7e-h).

These results suggest that the loss of *Tgfb1/2/3* using the αMHC-Cre strategy alters cardiomyocyte maturation before the P1 treatment window, within the embryonic heart, which is consistent with the profound changes in gene expression already observed at E17.5 (Fig. 3a).

### Analysis of fibroblast-cardiomyocyte communication in hearts of *Tgfb1/2/3* deleted mice

Our results suggested that cardiomyocyte-specific deletion of TGFβ impaired cardiac fibroblast differentiation in one of two possible ways. First, it is possible that defective fibroblast differentiation in the developing heart results in a less stiff and weakened ECM that fails to provide sufficient support or loading like conditions that cardiomyocytes need to properly differentiate [29, 30, 32, 33]. Second, it is also possible that defective fibroblast differentiation results in expression of inappropriate structural, matricellular and signaling genes, that together have a gain-of-function effect that pathologically alters cardiomyocyte maturation. To distinguish between these two mechanisms, we analyzed comparative effects in three different mouse models with defective fibroblast function or ECM composition. First, we performed RNA sequencing of hearts from *Tcf21^-/-^* mice versus *Tgfb1/2/3^fl/fl-αMHC-Cre^* mice at E17.5.

Deletion of *Tcf21* results in perinatal lethality, due to defective epicardial EMT and failure to form cardiac fibroblasts [20]. The data show almost no overlap in differentially expressed genes between hearts of *Tgfb1/2/3^fl/fl-αMHC-Cre^* and *Tcf21^-/-^* mice (1 shared upregulated gene, 7 shared downregulated genes) (Supplementary Fig. 8a,b). These results suggest that altered cardiac fibroblast function in hearts of *Tgfb1/2/3^fl/fl-αMHC-Cre^* mice, and not the loss of fibroblast activity, most likely underlies the phenotype observed. Moreover, analysis of hearts from *Col1a2/Col6a2* double null mice at 4 weeks of age that showed a similar defect in ECM stiffness with cardiomyopathy [41], failed to show the same altered gene expression profile as *Tgfb1/2/3^fl/fl-αMHC-Cre^* mice (Supplementary Fig. 8c-i). For example, the cardiomyocyte fetal gene *Tnni1* failed to be extinguished only in hearts from *Tgfb1/2/3^fl/fl-αMHC-Cre^* mice, nor were *Atp2a1*, *Ryr1*, *Tnni2*, and *Mylpf* skeletal muscle genes ectopically induced in *Col1a2/Col6a2* null hearts, nor was there continued upregulation of the fibroblast genes *Tgm2* and *Fbn2*, as observed in hearts from *Tgfb1/2/3^fl/fl-αMHC-Cre^* mice (Supplementary Fig. 8c-i). Finally, we also compared gene expression signatures of hearts from mice at P7 with developmental myocyte-specific deletion of *Itgb1*, which results in unloading of the myocytes and heart failure due to loss of integrin-ECM binding [43]. Hearts from *Itgb1^fl/fl-αMHC-Cre^* mice displayed an increase in expression of immature genes, such as *Cdkn1a*, like hearts from *Tgfb1/2/3^fl/fl-αMHC-Cre^* mice at P7; however, loss of *Itgb1* did not result in ectopic induction of skeletal muscle genes such as *RyR1* and *Mylpf* at P7 (Supplementary Fig. 8j-n). Taken together these results suggest that deletion of *Tgfb1/2/3* in cardiomyocytes of the embryonic heart leads to fibroblast reprogramming that leads to misdifferentiation of cardiomyocytes in the heart.

To examine whether TGFβ generated by cardiomyocytes in the developing heart was simply having an autocrine effect by signaling back to cardiomyocytes themselves to promote maturation, we deleted canonical downstream TGFβ signaling effectors *Smad2* and *Smad3*, or separately, *Tgfbr1 and Tgfbr2* from the cardiomyocyte using the same *αMHC-Cre* transgene.

At 6 weeks of age, *Smad2/3^fl/fl-αMHC-Cre^* mice did not display a reduction in cardiac function nor left ventricular dilation (Fig. 6a-b). Cardiomyocyte maturation assessed by quantifying nucleation at P21 also showed no effect in *Smad2/3^fl/fl-αMHC-Cre^* cardiomyocytes, unlike cardiomyocytes from hearts of *Tgfb1/2/3^fl/fl-αMHC-Cre^* mice (Fig. 6c). Because TGFβ could be regulating cardiomyocyte maturation through non-canonical TGFβ signaling pathways as well, we also deleted TGFβ receptor I and II, from the cardiomyocyte. Hearts from *Tgfbr1/2^fl/fl-αMHC-Cre^* mice also failed to show altered function or dilation in comparison to control littermates at 6 weeks of age (Fig. 6d,e), nor was there a maturation defect in percentage of mono-and bi-nucleated cardiomyocytes at P21 (Fig. 6f). Additionally, neither *Smad2/3^fl/fl-αMHC-Cre^* mice nor *Tgfbr1/2^fl/fl-αMHC-Cre^* displayed cardiac fibrosis as measured by picrosirius red staining and periostin staining (Fig. 6g), in comparison to abundant fibrosis in the hearts of *Tgfb1/2/3^fl/fl-αMHC-Cre^* mice at 6 weeks of age. Lastly, the loss of *Smad2/3* and *Tgfbr1/2* from cardiomyocytes did not result in ectopic expression of skeletal muscle-specific genes, *Atp2a1*, *Mylpf*, and *Tnni2* (Fig. 6h-j). Therefore, the defect in cardiomyocyte maturation in the hearts of *Tgfb1/2/3^fl/fl-αMHC-Cre^* mice and the resulting pathology is not due to failed autocrine TGFβ signaling back to the cardiomyocyte.

**Fig. 6.**
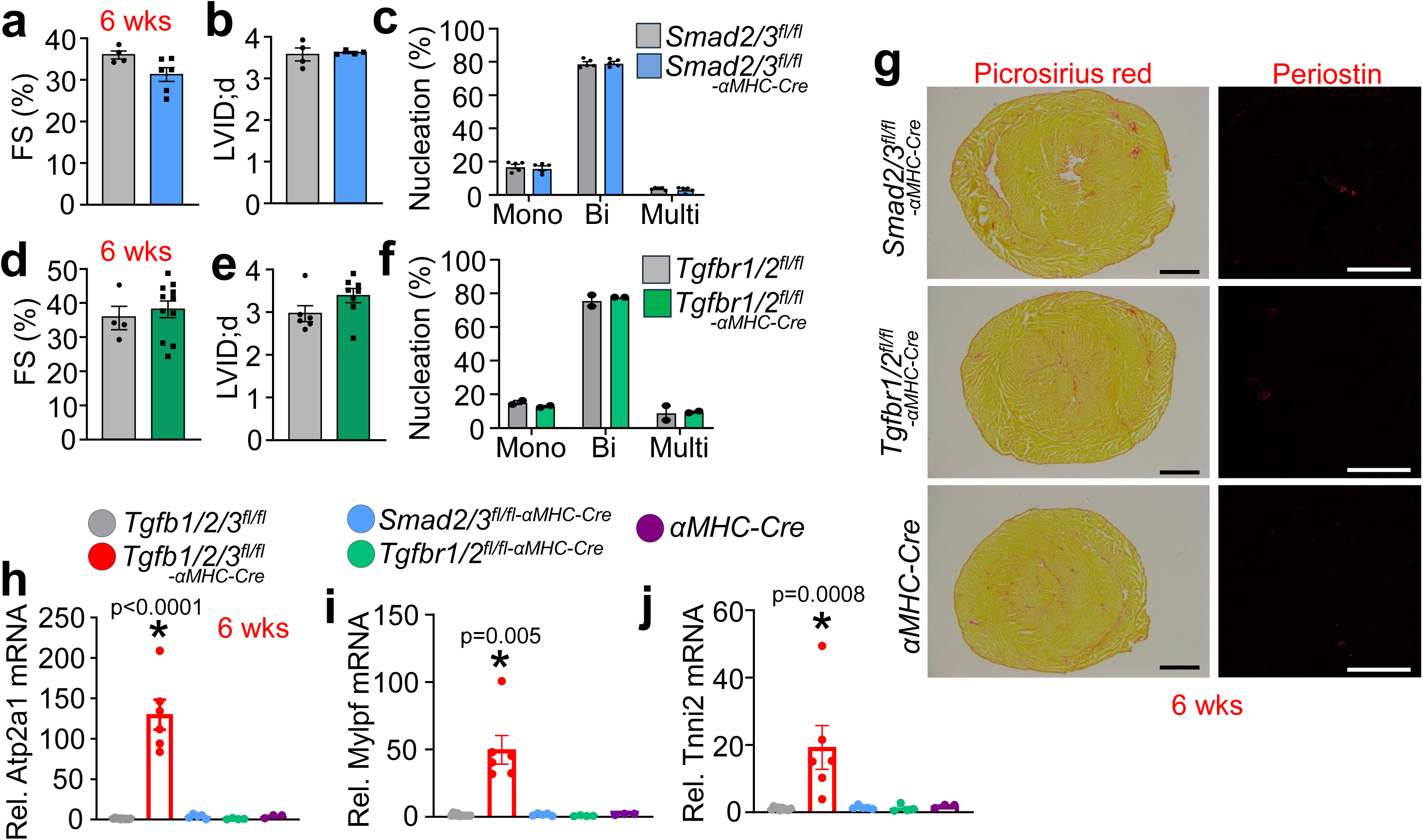
Cardiomyocyte-specific deletion of *Tgfbr1* and *Tgfbr2* or *Smad2* and *Smad3* does not impair cardiomyocyte maturation. **a**. Echocardiography measured fractional shortening (FS%) in the hearts of *Smad2/3^fl/fl-αMHC-Cre^* vs *Smad2/3^fl/fl^* mice at 6 weeks of age. n=4-6 mice per group. **b**. Echocardiography measured left ventricular internal diameter in diastole (LVID;d) in hearts of *Smad2/3^fl/fl-αMHC-Cre^* vs *Smad2/3^fl/fl^* mice at 6 weeks of age. n=4-6 mice per group. **c**. Quantification of mononucleation, binucleation, and multinucleation of isolated cardiomyocytes from 5 hearts each from *Smad2/3^fl/fl-αMHC-Cre^* vs *Smad2/3^fl/fl^* mice at P21. Minimally 200 cardiomyocytes were quantified per heart. **d**. Echocardiography measured fractional shortening (FS%) in the hearts of *Tgfbr1/2^fl/fl-αMHC-Cre^* vs *Tgfbr1/2^fl/fl^* mice at 6 weeks of age. n=4-11 mice per group. **e**. Echocardiography measured left ventricular internal diameter in diastole (LVID;d) in hearts *Tgfbr1/2^fl/fl-αMHC-Cre^* vs *Tgfbr1/2^fl/fl^* mice at 6 weeks of age. **f**. Quantification of mononucleation, binucleation, and multinucleation of isolated cardiomyocytes from two hearts each from *Tgfbr1/2^fl/fl-αMHC-Cre^* vs *Tgfbr1/2^fl/fl^* mice at P21. Minimally 200 cardiomyocytes were quantified per heart. **g**. Representative histological heart cross-sections with picrosirius red staining (left panel, scale bar: 1 mm) and periostin staining (red, right panel, scale bar: 100 µm) from hearts of *Smad2/3^fl/fl-αMHC-Cre^*, *Tgfbr1/2^fl/fl-αMHC-Cre^*, and *αMHC-Cre* mice at 6 weeks of age. **h-j**. Quantitative PCR analysis of skeletal muscle-specific genes *Atp2a1*, *Mylpf*, and *Tnni2* from hearts of mice with the genotypes shown at 6 weeks of age. Genes are standardized to 18S ribosomal RNA gene expression. n=3-10 mice per group. *p<0.05. a,b,d,e. Data are presented as mean +/-SEM. Student’s T-test was used to assess statistical significance. c,f. Data are presented as mean +/-SEM. Two-way ANOVA with a Tukey’s multiple comparison test was used to assess statistical significance. h. Data are presented as mean +/-SEM. One-way ANOVA with a Tukey’s multiple comparison test was used to assess statistical significance. I,j. Druskal-Wallis test with Dunn’s multiple comparisons test was performed to assess statistical significance of non-normally distributed data. *p<0.05

To further assess the signaling between cardiomyocytes and fibroblasts in the maturing heart we performed single nuclear sequencing of hearts from *Tgfb1/2/3^fl/fl-αMHC-Cre^* mice at P7, versus littermate controls. Analysis of the data showed no appearance of a unique cell population between hearts containing or lacking *Tgfb1/2/3* in cardiomyocytes versus control (Fig. 7a).

**Fig. 7.**
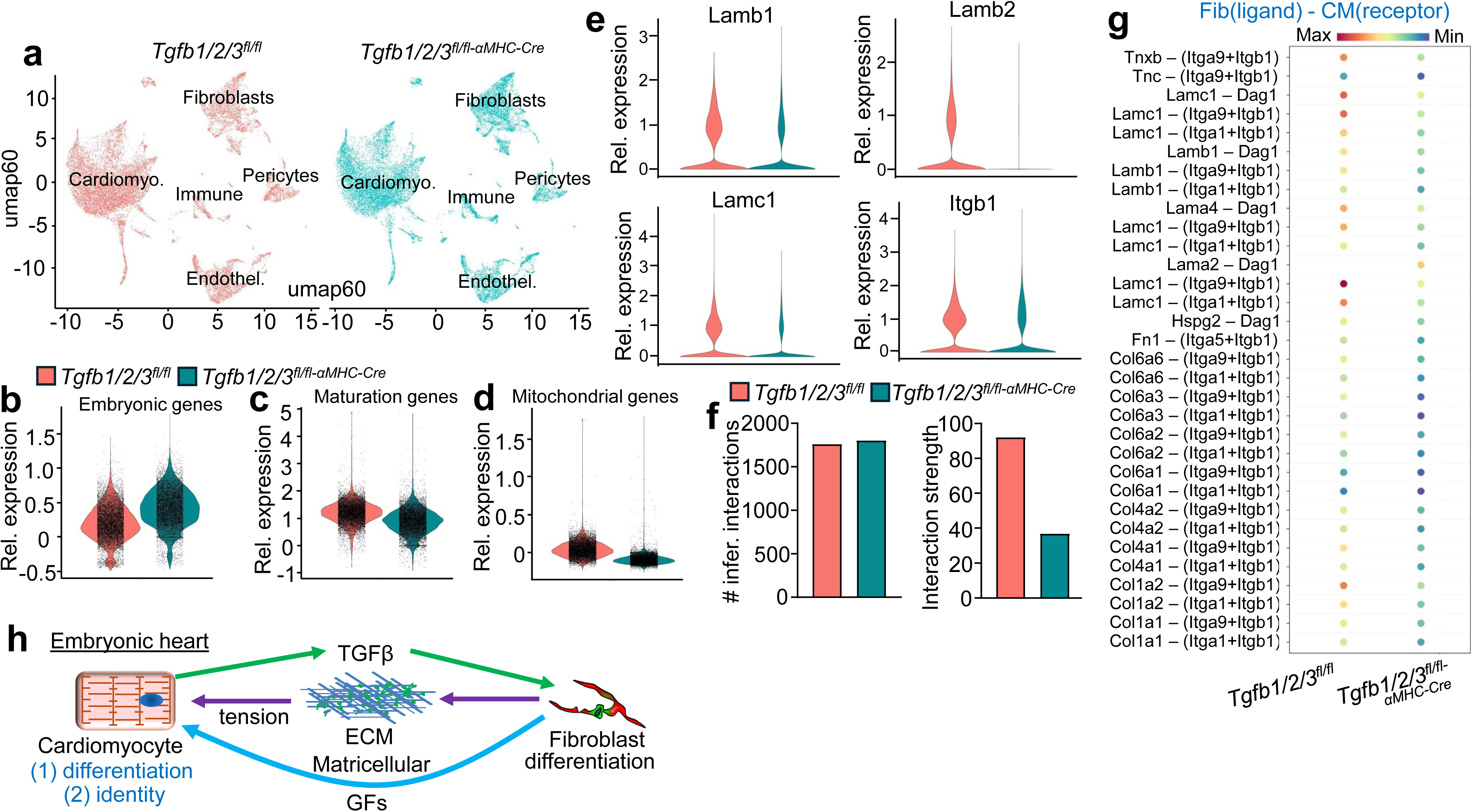
Single nuclear sequencing reveals reduced attachment complexes in hearts of *_Tgfb1/2/3fl/fl-αMHC-Cre_* _mice_. **a**. Uniform Manifold Approximation and Projections (UMAPs) displaying cellularity from hearts of *Tgfb1/2/3^fl/fl^* and *Tgfb1/2/3^fl/fl-αMHC-Cre^* mice at P7. **b**. Composite scoring of genes associated with embryonic cardiomyocytes based on expression of *Myh7*, *Tnni1*, *Myl4*, and *Myl7*. Each dot represents a nucleus scored based on enrichment in the embryonic cardiomyocyte gene signature. **c**. Composite scoring of relative expression of mature cardiomyocyte gene expression assessed by expression of *Myh6*, *Tnni3*, *Myl3*, and *Myl2*. Each dot represents a nucleus scored based on enrichment in the embryonic cardiomyocyte gene signature. **d**. Composite scoring of relative gene expression associated with mitochondrial maturation-based genes listed in Supplementary Table 3. **e**. Violin plot showing relative expression of *Lamb1*, *Lamb2*, *Lamc1*, and *Itgb1* in cardiomyocytes from *Tgfb1/2/3^fl/fl^* (red) and *Tgfb1/2/3^fl/fl-αMHC-Cre^* (blue). **f**. CellChat analysis of the number of inferred interactions and interaction strength between cardiac fibroblast ligands (cell-cell interactions or ECM-cell interactions) and cardiomyocyte receptors from hearts of *Tgfb1/2/3^fl/fl-αMHC-Cre^* mice and littermate controls. **g**. Dotplot of predicted differential ligand and receptor interactions between fibroblast (Fib) ligands paired with cardiomyocyte (CM) receptors. The scale located at the top of the figure displays strength of the predicted interaction. **h**. Schematic model of embryonic heart development where TGFβ generated by cardiomyocytes signals to tissue resident cardiac fibroblasts to initiate their differentiation and generation of proper ECM, matricellular, and growth factors (GF) that collectively signaling to maturing cardiomyocytes to ensure their differentiation and identity.

Given our previous data we scored all cardiomyocytes based on the expression of known genes associated with embryonic cardiomyocytes (*Myh7*, *Tnni1*, *Myl4*, *Myl7*) and found enrichment of embryonic genes in cardiomyocytes from *Tgfb1/2/3^fl/fl-αMHC-Cre^* hearts compared to control littermates (Fig. 7b), as well as reduced maturation markers (*Myh6*, *Tnni3*, *Myl3*, *Myl2*) (Fig. 7c). We also observed that several mitochondrial genes, which track with augmented maturation, were reduced in hearts from *Tgfb1/2/3^fl/fl-αMHC-Cre^* mice (Fig. 7d, Supplementary Table 3). Interestingly, cardiomyocytes from *Tgfb1/2/3^fl/fl-αMHC-Cre^* mice showed clear downregulation in laminins including *Lamb1*, *Lamb2*, and *Lamc1* as well as reduced expression of *Itgb1* (Fig. 7e). These data suggest impaired attachment complexes between cardiomyocytes and their respective basement membrane. Given the defect in fibroblast gene expression in hearts from *Tgfb1/2/3^fl/fl-αMHC-Cre^* mice (Fig. 2g-i) we used CellChat to focus on fibroblast ligands communicating with cardiomyocyte receptors (Fig. 7f,g). While the number of inferred fibroblast-cardiomyocyte interactions remained the same between hearts of *Tgfb1/2/3^fl/fl-αMHC-Cre^* mice and their control littermates, the interaction strength was reduced with loss of TGFβ, suggesting impaired signaling from the fibroblast (Fig. 7f). We identified a reduction in many basement membrane related interactions including laminins (*Lamc1*, *Lamb1*, *Lama4*, *Lama2*), perlecan, fibronectin, collagen VI, and collagen IV with cardiomyocyte integrin receptors (*Itga9*, *Itgb1*, *Itga1*, *Itga5*) (Fig. 7g). Therefore, the loss of cardiomyocyte maturation in hearts of *Tgfb1/2/3^fl/fl-αMHC-Cre^* mice is also associated with reduced basement membrane interactions and reduced fibroblast direct communication.

The ability of TGFβ to refine the ECM and provide additional signaling effects to augment myocyte maturation was also examined in mice using a gain-of-function approach that similarly impacted the early developmental period of the heart with a Mhy6 promoter driven transgene. We employed a latency escaping TGFβ1 mutant protein, which we described previously as generating fibrosis and cardiac hypertrophy at 6 months of age [64]. Interestingly, RNA sequencing analysis of hearts from control and *αMHC-Tgfb1* transgenic mice at 6 months of age showed increased expression of genes that would enhance ECM stability and cell-cell attachment, as well as genes that augment EMT such as *Mfap4*, *Lox*, *Ctgf*, *Col8a1*, *Nr4a1*, *Bgn*, *Itgbl1*, *Ncam1*, *Nrg1*, *Angptl7*, *Angptl4*, *Serpina3n*, *Adamtsl2*, *Sfrp2*, *Efemp1*, and *Neb* (Supplementary Table 4). Thus, TGFβ is critical for the developmental maturation of the cardiac fibroblast and the ECM it generates and refines, collectively programming cardiomyocyte maturation and proper cellular identity.

## DISCUSSION

Here we demonstrate that cardiomyocyte-specific deletion of TGFβ ligand encoding genes, *Tgfb1*, *Tgfb2*, and *Tgfb3*, results in cardiac fibrosis by 6 weeks of age and lethal heart failure by ∼6 months of age. Assessment of ECM organization in response to acute injury revealed that collagen fibers in the hearts of *Tgfb1/2/3^fl/fl-αMHC-Cre^* hearts were shorter and disorganized in comparison to control hearts, suggesting a disease mechanism related to cardiac fibroblasts. The embryonic cardiac fibroblast generates an ECM that is rich in fibronectin, fibrillin-2, loxl2, transglutaminase 2, and periostin, which also promotes cardiomyocyte cell cycle activity in the early heart [27–29, 31]. The postnatal heart shows replacement of fetal ECM components with fibrillar collagens and laminins that can drive cardiomyocyte maturation through increasing stiffness [27, 29, 75]. Single nuclear RNA sequencing of hearts from *Tgfb1/2/3^fl/fl-αMHC-Cre^* mice at P7 revealed that cardiac fibroblasts also failed to upregulate genes involved in postnatal ECM maturation including *Lama2*, *Lama4*, *Lamb1*, *Lamb2*, and *Lamc1*. By comparison, Laminin-511 (α5β1γ1) and Laminin-521 (α5β2γ1) were identified in a screen of ECM proteins that enhanced maturation of pluripotent stem cell-derived cardiomyocytes [76]. In addition to reduced expression of laminins in fibroblasts, cardiomyocytes from *Tgfb1/2/3^fl/fl-αMHC-Cre^* hearts at P7 showed downregulation of integrins and other attachment proteins, collectively leading to reduced load sensing conditions that would lead to defective cardiomyocyte maturation.

TGFβ is traditionally assumed to be released by immune cells, fibroblasts and endothelial cells, although here we observed a requirement for TGFβ expression in cardiomyocytes of the early developing heart. This observation fits with the known structural design of TGFβ1/2/3 proteins, which contain their own extracellular linkage latency-associated peptide (LAP) domain. The LAP domain sequesters the co-translated TGFβ ligand portion and renders its release sensitive by stretch through direct binding to integrins, or by proteolytic activity regulated by other signals. However, the TGFβ-LAP complex with latent TGFβ binding protein (LTBP) and fibrillin is not easily assembled and shuttled throughout the cardiac ECM. Thus its production by cardiomyocytes represents an outward signal at the level of the basement membrane to then program fibroblasts within the ECM. Indeed, mice with a germline mutation in the RGD motif of the LAP domain of *Tgfb1*, which cannot undergo stretch-mediated activation, result in the same deleterious phenotype as *Tgfb1* germline null mice [77]. Our results suggest that the purpose of this TGFβ signal is not autocrine, but instead specific to fibroblasts given the observation that cardiomyocyte-specific deletion of TGFβ receptors I and II (*Tgfbr1/2^fl/fl-αMHC-Cre^*) or canonical TGFβ signaling proteins, Smad2/3 (*Smad2/3^fl/fl-αMHC-Cre^*) did not produce the phenotypic effects associated with *Tgfb1/2/3* deletion using the same αMHC-Cre transgene.

The cardiomyocyte-specific loss of TGFβ ligands from the heart not only reduced expression of maturation genes in the heart, but also unexpectedly resulted in ectopic induction of multiple skeletal muscle-specific genes (Fig. 7h). To our knowledge, evidence of skeletal muscle specific-gene expression in the heart has only been observed in four previous studies in mice, all of which involved cardiomyocyte-specific deletion of global chromatin and epigenetic regulators; *Hdac1/2*, *Ezh2*, *Jarid2*, and *Chd4* (Supplementary Table 2, [70–73]). Indeed, these cardiac specific gene deleted mice also showed cardiomyopathy in late neonatal or early adulthood, similar to the phenotype of *Tgfb1/2/3^fl/fl-αMHC-Cre^* mice. This ectopic induction of a selective skeletal muscle gene program could easily disrupt the finely tuned physiology of the heart leading to cardiomyopathy and early adulthood lethality. Interestingly, single nuclear RNA sequencing performed on hearts of *Tgfb1/2/3^fl/fl-αMHC-Cre^* mice at P7 revealed downregulation of numerous epigenetic regulators including *Hdac1*, *Hdac3*, *Hdac10*, *Ezh2*, and *Cdk8* (FC<-1.25, p<0.05). This ectopic skeletal muscle gene expression program with *Tgfb1/2/3* deletion began at E17.5 with induction of *Ryr1* and *Casq1*, then by P7 *Mylpf* induction was also observed, followed by full induction a wide array of skeletal muscle genes by 4 weeks of age. This gradual transcriptional induction suggests the continued influence of an altered ECM environment.

The reduced stiffness of the ECM due to deletion of *Tgfb1/2/3* from cardiomyocytes clearly leads to a defect in myocyte maturation, as discussed above. However, multiple lines of evidence suggest that reduced ECM stiffness alone should not alter the identity of maturing cardiomyocytes towards the skeletal muscle gene program, implicating a more profound pathological effect. For example, hearts from germline *Col1a2^-/-^* mice that have a prominent and singular defect in ECM stiffness and induction of the fibrotic response did not show inappropriate cardiomyocyte differentiation [41], and Ullrich muscular dystrophy patients with mutations in collagen 6 genes show defects in skeletal muscle maturation due to loss of basement membrane coupling [78]. To produce an even greater defect in the basement membrane and ECM support network of the heart we generated *Col1a2/Col6a2* double null mice that showed cardiomyopathy by 4 weeks of age but failed to show inappropriate cardiomyocyte differentiation compared to 4-week-old hearts from *Tgfb1/2/3^fl/fl-αMHC-Cre^* mice (Supplementary Fig. 8 c-i). As yet another line of evidence we also disrupted the *Itgb1* gene in cardiac myocytes, given the observation that blocking the interaction of integrin β1 between postnatal cardiomyocytes and the ECM impaired cardiomyocyte maturation [31], and cardiomyocytes from *Tgfb1/2/3^fl/fl-αMHC-Cre^* mice display a downregulation of integrin β1 at P7. Cardiomyocyte-specific deletion of *Itgb1* using MLC2v-Cre also resulted in fibrotic dilated cardiomyopathy at 6 months of age resembling *Tgfb1/2/3^fl/fl-αMHC-Cre^* mice ([79], Supplementary Fig. 1). Here, early developmental deletion of *Itgb1* from cardiomyocytes with the same αMHC-Cre transgene, which unloads the myocyte from the ECM, did not negatively affect cardiomyocyte differentiation at P7, as observed in hearts from *Tgfb1/2/3^fl/fl-αMHC-Cre^* mice (Supplementary Fig. 8j-n). Lastly, while hearts from *Tgfb1/2/3^fl/fl-αMHC-Cre^* mice at E17.5 already showed a profile of reduced maturation and ectopic skeletal muscle gene expression,*Tcf21^-/-^* mice at E17.5 that lack fibroblasts altogether did not (Supplementary Fig. 4a-c, Supplementary Fig. 8a-b), suggesting that the primary mechanism is not due to a lack of ECM support and cardiomyocyte load sensing (Fig. 7h).

Our collective data discussed above point towards an alternate explanation, whereby defective fibroblast differentiation in the early heart of *Tgfb1/2/3^fl/fl-αMHC-Cre^* mice leads to a pathologically altered ECM and matricellular network that sends an inappropriate signal to co-maturing cardiomyocytes. For example, hearts from *Tgfb1/2/3^fl/fl-αMHC-Cre^* mice at P7 showed persistent expression of a fetal or the injury ECM program including fibronectin, periostin, transglutaminase 2, lumican and tenascin C. Indeed, fibronectin was previously shown to uniquely drive cardiomyocyte proliferation at later time points [19]. Even more provocatively, fibroblasts from adult hearts of *Tgfb1/2/3^fl/fl-αMHC-Cre^* mice expressed several unique regulatory genes not typically identified in mouse models of heart disease and fibrosis [66, 67]. Supplementary Table 1 describes 43 of these unorthodox fibroblast genes, which are more typically observed in cancer associated fibroblasts, such as *Bdnf, Ereg, Cst6, Bgn, Sfrp1, Mme1, Gpc3, Amigo2*, and *Olfml2a*. Genes associated with mesenchymal cell proliferation and function were also differentially expressed, such as *Wisp2, Gdf6, Hes1, Pttg1, Asb2, Meis3,* and *IL-18*. Indeed, germline deletion of the *Tgfb1* gene results in postnatal lethality with a noncompact myocardium and fibrosis with signs of myocyte proliferation and necrosis at P7 and cardiomyocytes from these hearts appear to have inappropriate upregulation of major histocompatibility class II molecules (ectopic), which we also identified in hearts of adult *Tgfb1/2/3^fl/fl-αMHC-Cre^* mice [13].

Another interesting observation was that the expression of *Tgfb1*, but not *Tgfb2* or *Tgfb3*, was essential for cardiomyocyte maturation and thus, preservation of cardiac function. Much of our analysis was performed with the triple null mice because both *Tgfb2* and *Tgfb3* are made by cardiomyocytes in the early heart and the triple null strategy protected against potential compensation from *Tgfb2* and *Tgfb3*. However, it is possible that the three different TGFβ encoding genes are non-redundant, or that *Tgfb1* is uniquely expressed at a more critical time for proper heart maturation, or that the potency and levels of expression are dominated by *Tgfb1*. We also unsuccessfully attempted to rescue myocyte maturation and heart function in *Tgfb1/2/3^fl/fl-αMHC-Cre^* mice by administering MyoAAV-*Tgfb1* or rTGFβ1 to early neonates, suggesting that cardiomyocyte generated TGFβ programs heart maturation before birth. However, the lack of TGFβ generated by cardiomyocytes did not impact epicardial EMT nor EPDC migration, suggesting that fibroblasts were still generated and distributed throughout the heart, even if not properly differentiated.

TGFβ can also impact the immune response and possibly activity of tissue resident immune cells in the heart [80], although we did not observe a difference in the number of tissue resident immune cells or an induction of inflammatory cells in the hearts of *Tgfb1/2/3^fl/fl-αMHC-Cre^* mice at P7 (Supplementary Fig. 9). In the future it will be important to provide even more direct evidence that fibroblasts are the critical intermediary cell type that drives TGFβ-dependent maturation by deleting the genes encoding the two TGFβ receptors or Smad2/3 early in embryonic fibroblasts after epicardial EMT and EPDC migration. However, we currently lack the appropriate genetic tools to permit this more direct analysis in embryonic heart fibroblasts. Despite this qualification, the current study employed multiple independent lines of investigation to support the conclusion that delayed and inappropriate cardiomyocyte maturation in hearts of *Tgfb1/2/3^fl/fl-αMHC-Cre^* mice is mediated by defective cardiac fibroblast differentiation in the heart, which not only matures the ECM, but it also provides the appropriate environment to ensure proper cardiomyocyte identity (Fig. 7h).

## Acknowledgements

This work was supported by the National Institutes of Health grants to D.P.M (R01AR068286, R01AG082697), J.D,M (1R01HL160765), M.D.T (R01HL144067), K.C.H (Cancer Center Support Grant (P30CA046934) for the Mass Spectrometry Proteomics Shared Resource Facility (RRID: SCR_021988)), M.T.W (P30AR070549), M.A. (R01HL126705, R21ES037105, R41 HL172481) and an ARC grant from the Cincinnati Children’s Hospital Medical Center (Award #53632).

## Data and code availability

RNA-seq, Affymetrix array data and snRNA-seq datasets described in this paper are located in the GEO repository under accession number xxxxxx. The original code for bioinformatic analysis of the single nuclear sequencing results is uploaded to GitHub [https://github.com/cswoboda/NF-Myonuclei]. All other raw data matched to the figures and tables are available upon reasonable request.

