## Supplemental figures and tables for "Cardiomyocyte-expressed TGFβ signals to fibroblasts to program early heart maturation and adult myocyte identity"

#### Supplementary Figures 1 - 9

#### Supplementary Tables 1 - 4

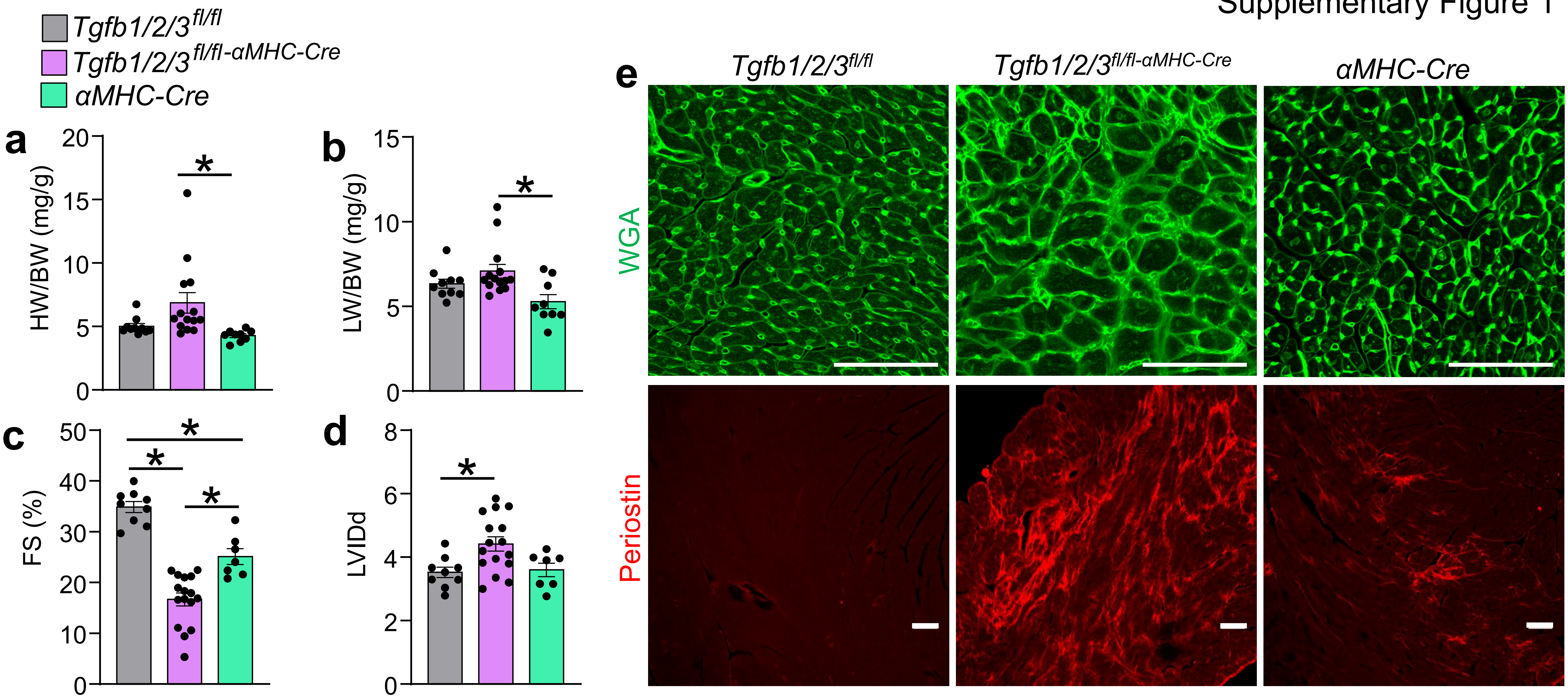


**Supplementary Fig. 1. Deletion of *Tgfb1/2/3* genes in cardiomyocytes *in vivo* results in progressive cardiac dysfunction and fibrosis at 6 months of age**

a) Heart weight to body weight ratio (HW/BW) in the indicated 3 genotypes of mice at 6 months of age. n=9-14 mice per group. *p<0.05.

b) Lung weight to body weight ratio (LW/BW) in the indicated 3 genotypes of mice at 6 months of age. n=9-14 mice per group. *p<0.05.

c) Fractional shortening (FS%) in the indicated 3 genotypes of mice at 6 months of age. n=7-16 mice per group. *p<0.05.

d) Left ventricular internal diameter in diastole (LVIDd) in the indicated 3 genotypes of mice at 6 months of age. n=7-16 mice per group. *p<0.05.

e) Representative images of immunostaining of histological heart sections for wheat germ agglutinin (WGA) (green, upper panel, scale bar: 100 μm) and periostin (red, bottom panel, scale bar: 200 μm) for indicated genotypes at 6 months of age. A-D. Data presented as mean +/- SEM. One-way ANOVAs used for statistical analysis.

### Supplementary Figure 2

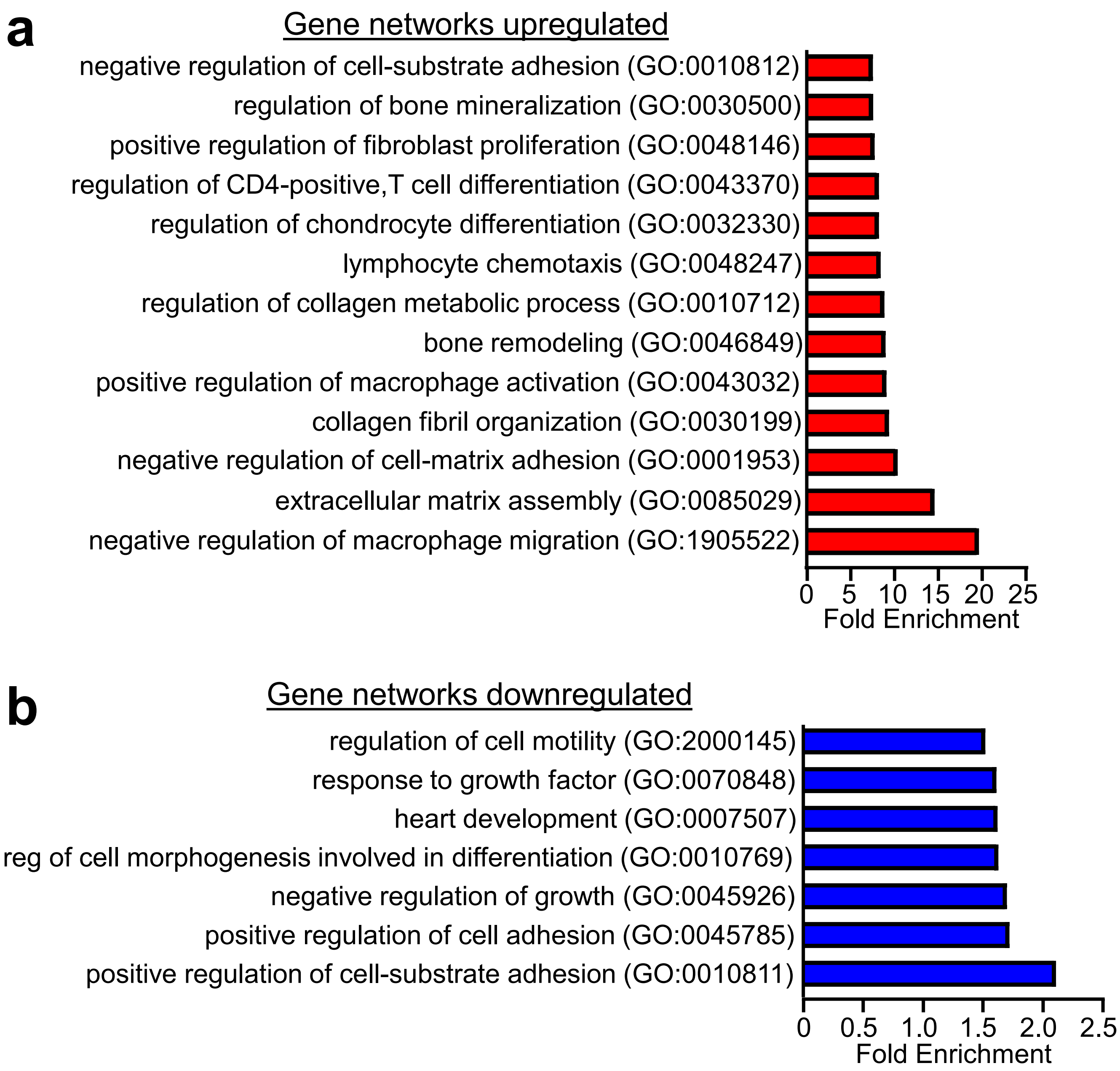


**Supplementary Fig. 2. Gene Ontology pathways from RNA sequencing of fibroblasts isolated from adult hearts of *Tgfb1/2/3*$^{fl/fl\text{-}\alpha MHC\text{-}Cre}$ versus *Tgfb1/2/3*$^{fl/fl}$ control mice.**
a,b. Gene Ontology (GO) pathways representing genes upregulated (a) and downregulated (b) in fibroblasts isolated from hearts of *Tgfb1/2/3*$^{fl/fl\text{-}\alpha MHC\text{-}Cre}$ mice versus *Tgfb1/2/3*$^{fl/fl}$ mice at 6 weeks of age. Cutoffs for GO analysis were genes with fold change >1.5 for upregulated genes or <-1.5 for downregulated genes and significance was analyzed with Panther and a Fisher Test with correction FDR <0.05.

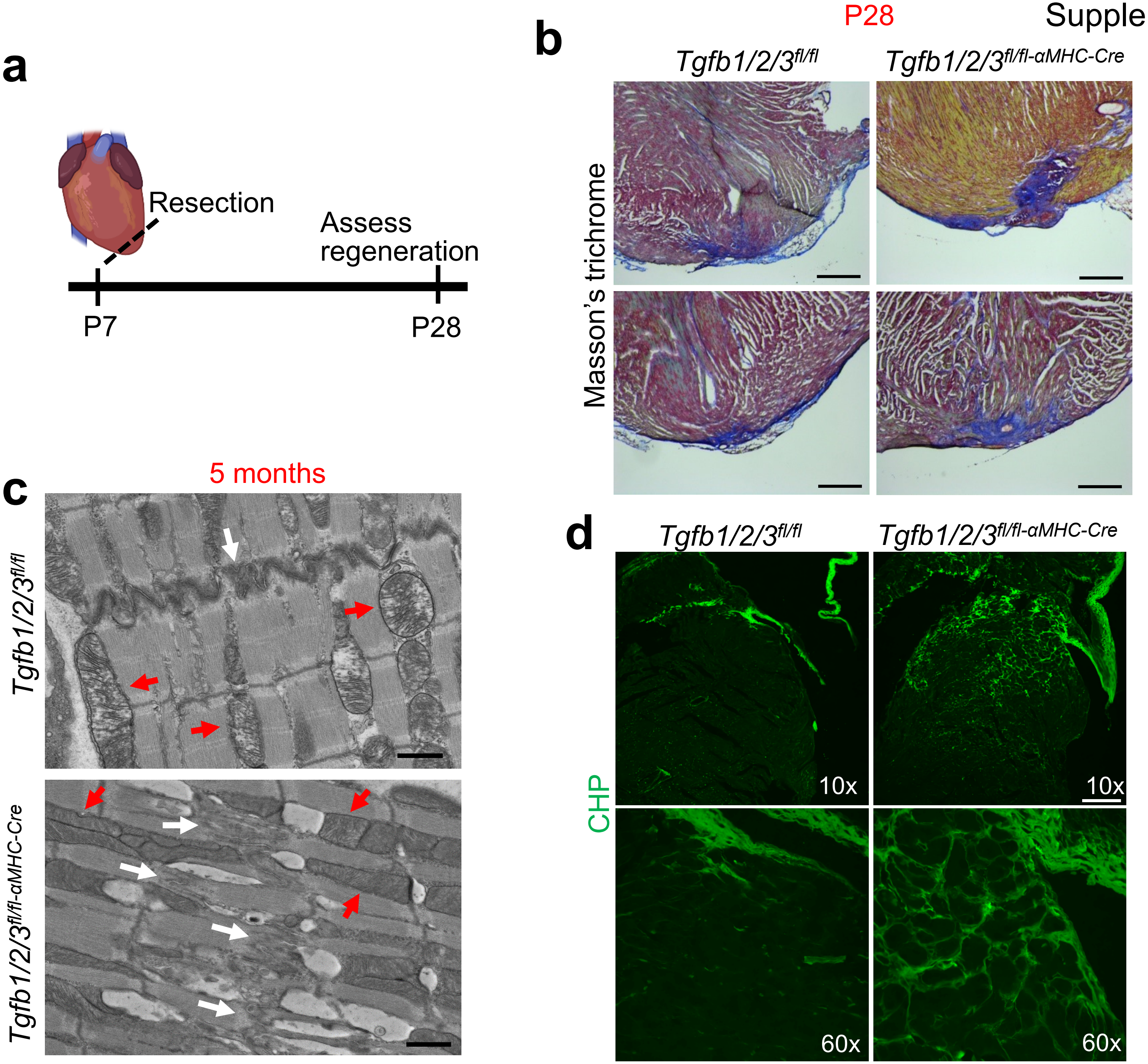
a
Resection
Assess regeneration
P7
P28
b
P28
Tgfb1/2/3fl/fl
Tgfb1/2/3fl/fl-αMHC-Cre
Masson's trichrome
c
5 months
Tgfb1/2/3fl/fl
Tgfb1/2/3fl/fl-αMHC-Cre
d
Tgfb1/2/3fl/fl
Tgfb1/2/3fl/fl-αMHC-Cre
CHP
10x
10x
60x
60x

**Supplementary Fig. 3. Hearts of *Tgfb1/2/3*$^{fl/fl\text{-}\alpha MHC\text{-}Cre}$ mice lack regenerative capacity, display ultrastructural changes, and exhibit improper collagen processing**

a) Schematic of the apical heart resection protocol in P7 mice. Approximately 15% of the apex of the heart was resected. Mice were sacrificed at P28 and assessed for potential regrowth of cardiac tissue or scar formation.

b) Masson's Trichrome histological heart staining of fibrosis (blue) from mice with P7 heart resection and analysis 21 days later in the 2 genotypes of mice shown. Scale bar is 0.5 mm

c) Representative transmission electron microscopy images from hearts of *Tgfb1/2/3*$^{fl/fl\text{-}\alpha MHC\text{-}Cre}$ mice versus *Tgfb1/2/3*$^{fl/fl}$ control mice at 5 months of age. White arrows indicate intercalated discs. Red arrows point to mitochondria. Scale bar: 1 μm.

d) Representative immunostaining for collagen hybridizing peptide (CHP, green) in histological sections from hearts of mice of the indicated genotypes at both 10X (upper panel) or 60X (bottom panel) objectives of the 2 indicated genotypes at 4 weeks of age.

Supplementary Figure 4

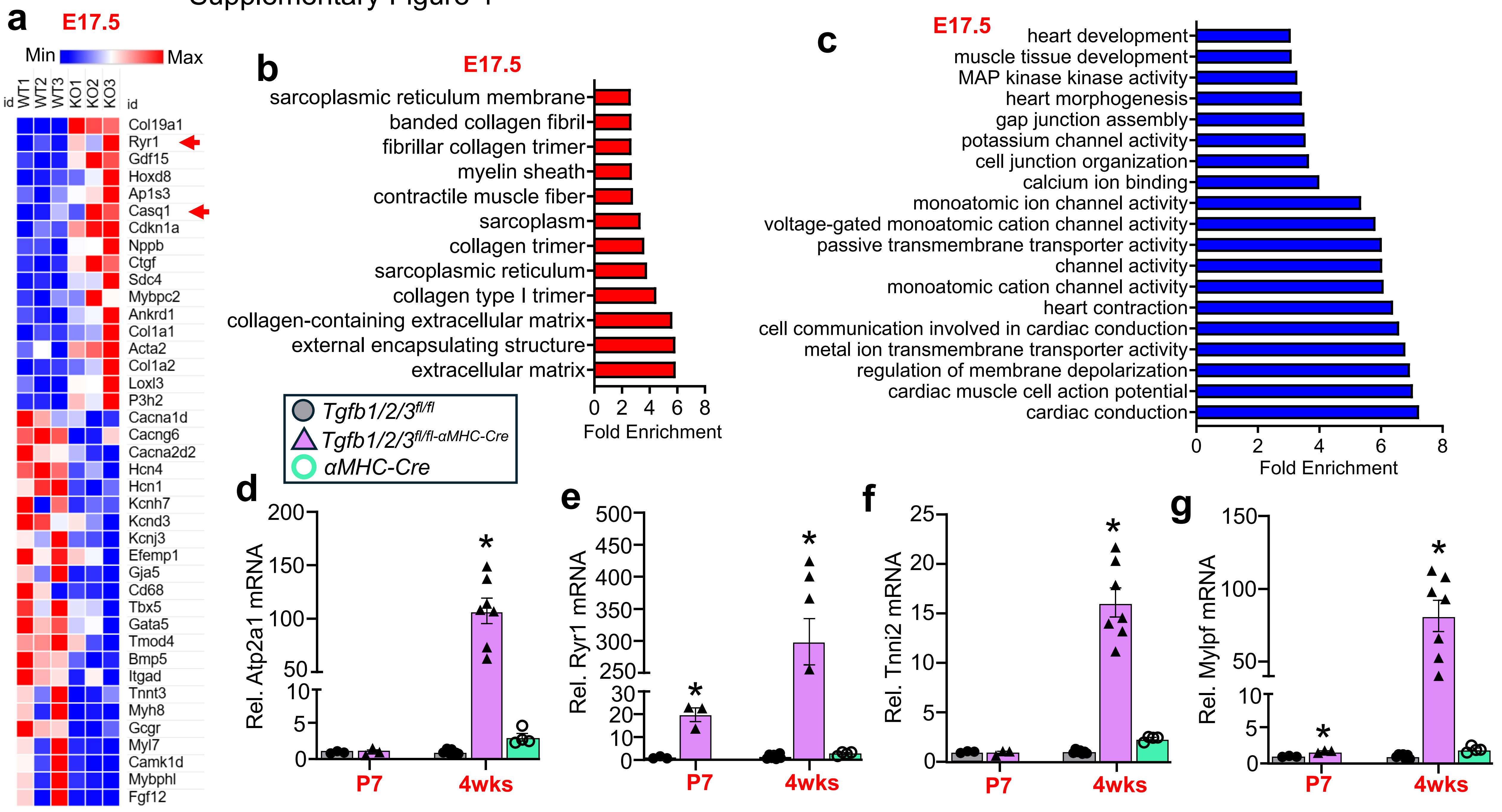

**Supplementary Fig. 4. Cardiomyocytes from *Tgfb1/2/3*$^{fl/fl\text{-}\alpha MHC\text{-}Cre}$ mice display impaired maturation at E17.5 and progressive ectopic skeletal muscle gene expression from P7 to 4 weeks of age.**

a) Heat map of select genes upregulated and downregulated in hearts of *Tgfb1/2/3*$^{fl/fl\text{-}\alpha MHC\text{-}Cre}$ mice versus wildtype (WT) control mice taken at E17.5 of gestation. Upregulated genes include some skeletal muscle-specific genes including *Ryr1*, *Casq1*, and *Ankrd1*. Downregulated genes include expression of several ion channels. Differential gene regulation was defined by FC>1.5 (upregulated) or FC<-1.5 (downregulated) with significance achieved at p<0.05

b) Gene Ontology pathway analysis of genes upregulated from hearts of mice described in a. Data were analyzed by ToppFun with a Probability Density Function Test and the correction was FDR <0.05.

c) Gene Ontology pathway analysis of genes downregulated from hearts of mice described in a. Data were analyzed by ToppFun with a Probability Density Function Test and the correction was FDR <0.05.

d-g) Quantitative PCR analysis of *Atp2a1, Ryr1, Tnni2,* and *Mylpf* at both P7 and 4 weeks of age from hearts of *Tgfb1/2/3*$^{fl/fl}$, *Tgfb1/2/3*$^{fl/fl\text{-}\alpha MHC\text{-}Cre}$, and αMHC-Cre mice. Relative expression was normalized to 18S ribosomal RNA expression. n=3-7 mice per group. *p<0.05. Data presented as mean +/- SEM. Two-Way ANOVAs were performed for statistical analysis.

Supplementary Figure 5

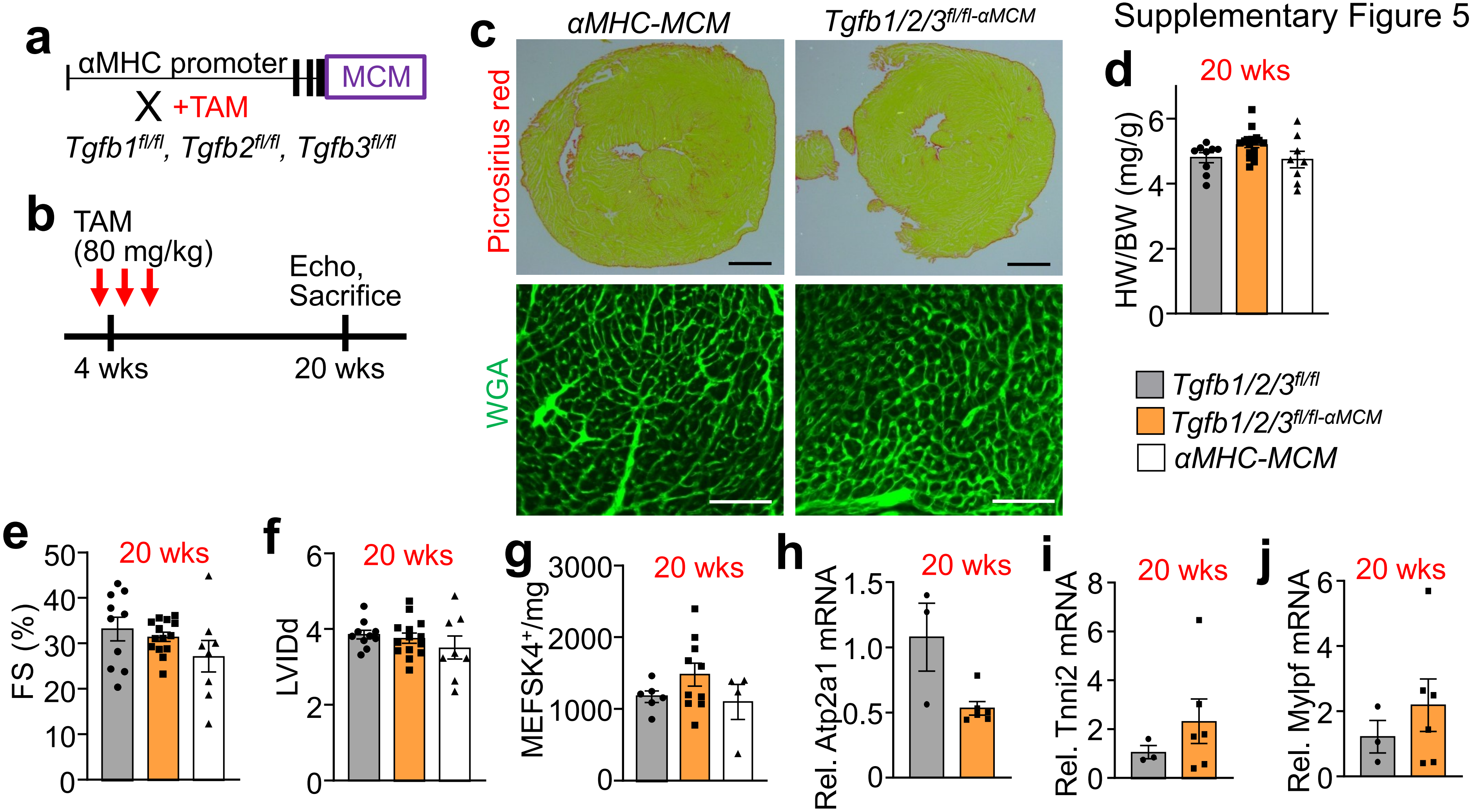

**Supplementary Fig. 5. Deletion of *Tgfb1/2/3* genes from myocytes of the adult heart does not alter cardiac function nor cardiomyocyte maturation**

a) Schematic showing the breeding strategy to generate *Tgfb1/2/3*$^{fl/fl\text{-}\alpha MCM}$ mice using the tamoxifen (TAM) inducible and cardiac specific transgene αMHC-MerCreMer (MCM).

b) Schematic showing the treatment strategy to delete *Tgfb1/2/3* genes in *Tgfb1/2/3*$^{fl/fl\text{-}\alpha MCM}$ mice over the course of 5 days with 80 mg/kg tamoxifen injection at 4 weeks of age. Mice were subject to echocardiography and sacrificed 16 weeks later at 20 weeks of age and processed.

c) Representative heart histological images of immuno-stained with Picrosirius red (upper panel, scale bar 1 mm) and WGA (lower panel, green, scale bar: 100 µm) from the 2 indicated genotypes of mice with adult deletion of *Tgfb1/2/3*.

d) Heart weight to body weight ratio (HW/BW) at 20 weeks of age in the 3 genotypes of mice used and processed at shown in a,b. n=8-17 mice per group.

e) Echocardiography measured fractional shortening (FS%) at 20 weeks of age in the 3 genotypes of mice used and processed at shown in a,b. n=8-14 mice per group.

f) Echocardiography measured left ventricular internal diameter in diastole (LVIDd) at 20 weeks of age in the 3 genotypes of mice used and processed at shown in a,b. n=8-14 mice per group.

g) Quantification of MEFSK4+ fibroblasts by flow cytometry from the left ventricle of mice from the indicated genotypes of mice at 20 weeks of age that were used and processed as shown in a, b. n=4-10 mice per group.

H-J. Quantitative PCR analysis of skeletal muscle specific genes, *Atp2a1*, *Mylpf*, and *Tnni2* from hearts of indicated genotypes at 20 weeks of age that were used and processed as shown in a,b. Gene expression was normalized to 18S ribosomal RNA expression. n=3-6 mice per group.

d-g. Data presented as mean +/- SEM. Statistical analysis was performed using One-way ANOVAs.
h-j. Data presented as mean +/- SEM. Statistical analysis was performed using Student’s T-tests.

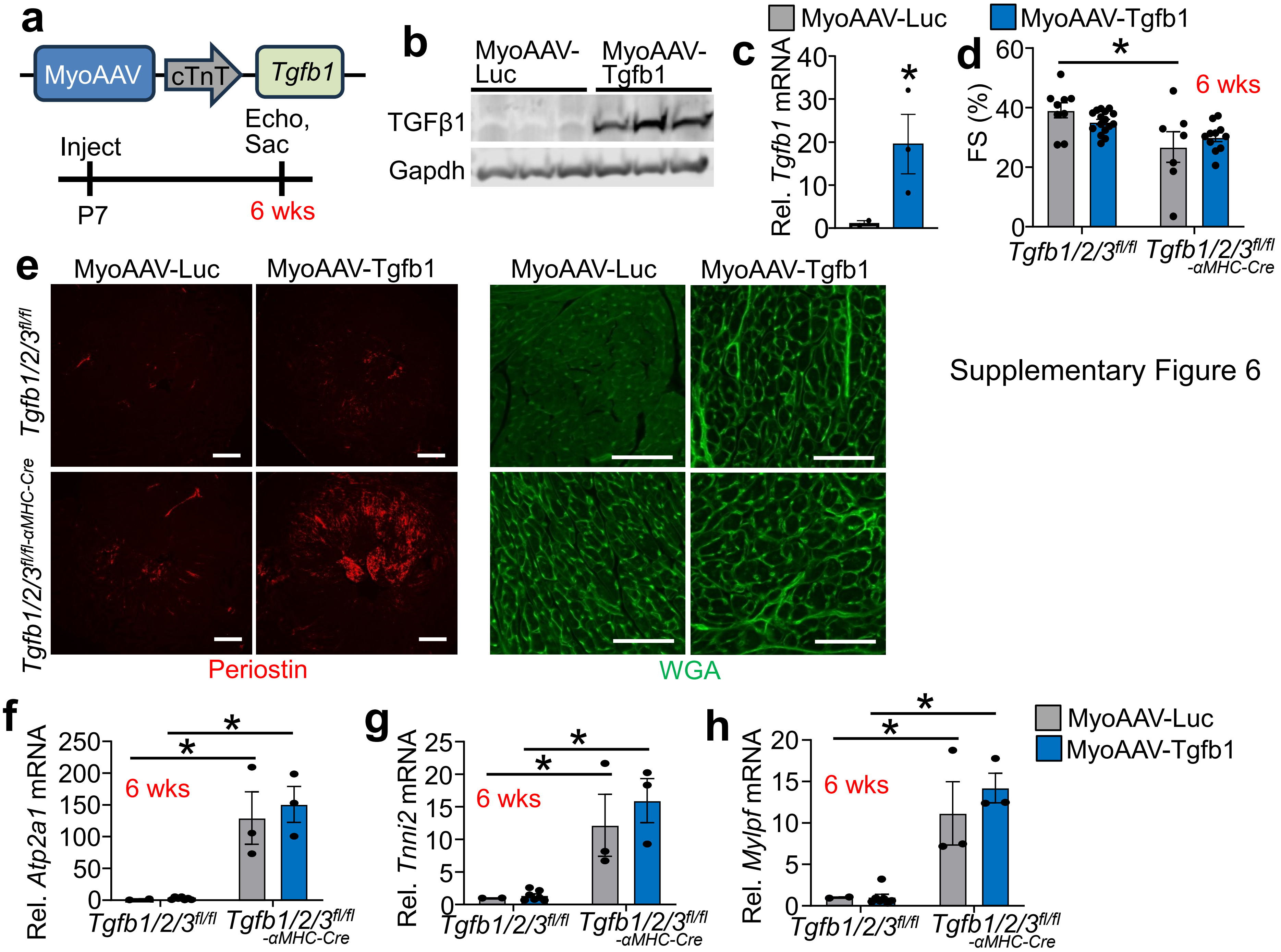


Supplementary Figure 6

**Supplementary Fig. 6. Postnatal Tgfb1 overexpression by MyoAAV does not rescue cardiac function nor cardiomyocyte maturation in *Tgfb1/2/3$^{fl/fl\text{-}\alpha MHC\text{-}Cre}$* mice**

a) Schematic of the MyoAAV-Tgfb1 design and experimental set up. Neonatal pups from *Tgfb1/2/3$^{fl/fl\text{-}\alpha MHC\text{-}Cre}$* and *Tgfb1/2/3$^{fl/fl}$* mice at P7 were injected with either a MyoAAV expressing *Tgfb1* under a cardiac troponin T promoter (MyoAAV-Tgfb1) or control MyoAAV expressing luciferase under a cardiac troponin T promoter (MyoAAV-Luc). Mice were then analyzed at 6 weeks of age.

b) Western blot analysis of TGFβ1 expression from hearts of *Tgfb1/2/3$^{fl/fl}$* mice treated with either MyoAAV-Luc or MyoAAV-Tgfb1 at P7 and sacrificed at 6 weeks of age. Gapdh was a tissue processing and western loading control.

c) Quantitative PCR analysis of *Tgfb1* overexpression from hearts of *Tgfb1/2/3$^{fl/fl}$* mice treated with either MyoAAV-Luc or MyoAAV-Tgfb1. *Tgfb1* expression was normalized to 18S ribosomal RNA expression. n=2-3 mice per group. $^*p<0.05$.

d) Echocardiography measured fractional shortening (FS%) at 6 weeks of age in the 2 genotypes of mice injected at P7 with either of the 2 viruses and analyzed at 6 weeks of age. n=7-16 mice per group. $^*p<0.05$.

e) Representative cardiac histological images of periostin (red) immunofluorescent staining (left panels, scale bar: 500 µm) and WGA staining (right panels, green, scale bar: 100 µm). Histological sections are from hearts of *Tgfb1/2/3$^{fl/fl\text{-}\alpha MHC\text{-}Cre}$* mice or *Tgfb1/2/3$^{fl/fl}$* controls treated with either MyoAAV-Luc or MyoAAV-Tgfb1.

f-h) Quantitative PCR analysis of skeletal muscle-specific genes from hearts of *Tgfb1/2/3$^{fl/fl\text{-}\alpha MHC\text{-}Cre}$* or control *Tgfb1/2/3$^{fl/fl}$* mice treated with either MyoAAV-Luc or MyoAAV-Tgfb1. Gene expression was normalized to 18S ribosomal RNA expression. n=2-7 mice per group. $^*p<0.05$.

Data are displayed as mean +/- SEM. Statistical analysis was performed using Student's T-test (c) or using a Two-way ANOVA (d, f-h).

### Supplementary Figure 7

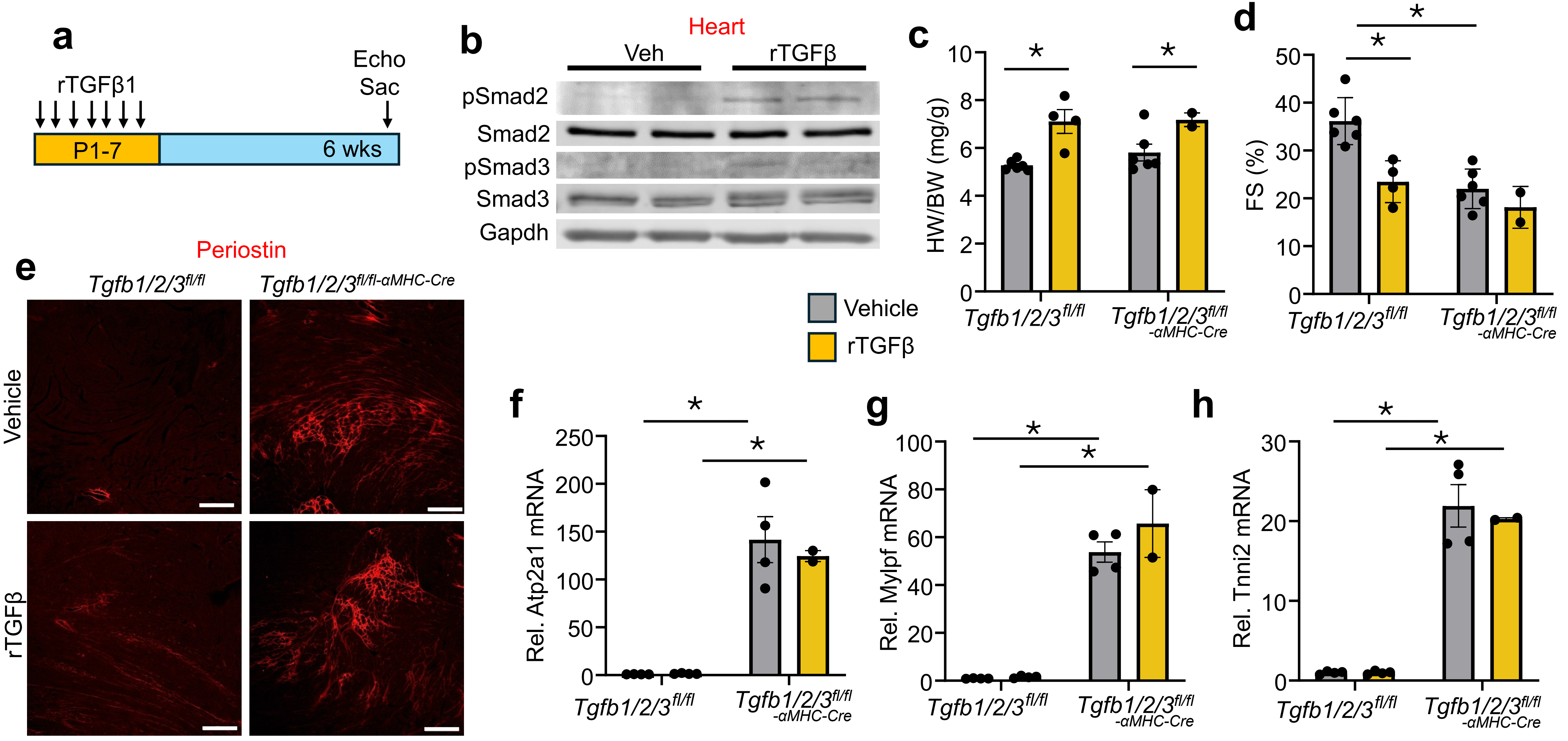

**Supplementary fig. 7. Administration of rTGFβ1 at P1-P7 does not rescue altered cardiac function or cardiomyocyte maturation of hearts from *Tgfb1/2/3*$^{fl/fl\text{-}\alpha MHC\text{-}Cre}$ Mice.**

a) Schematic of the dosing scheme used. *Tgfb1/2/3*$^{fl/fl\text{-}\alpha MHC\text{-}Cre}$ mice or littermate controls *Tgfb1/2/3*$^{fl/fl}$, were injected i.p. with 0.5 µg rTGFβ1 daily from P1-P4 and 1 µg rTGFβ1 daily from P5-P7 or equal volume vehicle control.

b) Western blot analysis from heart to assess phosphorylation of SMAD2 and SMAD3 following treatment with rTGFβ1. Wild-type pups at P3 were injected i.p. with 1 µg of rTGFβ1 and sacrificed 2 hours later. Hearts were homogenized and protein extracts used to measure total and phosphorylated SMAD2 and SMAD3. Gapdh was used as a protein extract and loading control.

c) Heart-weight to body-weight ratio (HW/BW) of mice from the 2 genotypes of mice treated with drug or vehicle as shown and sacrificed at 6 weeks of age. n=2-6 mice per group. *$p<0.05$.

d) Echocardiography measured fractional shortening (FS%) analysis mice from the 2 genotypes of mice treated with drug or vehicle as shown and processed at 6 weeks of age. n=2-6 mice per group. *$p<0.05$.

e) Representative heart histological images with immunofluorescent staining for periostin from the 2 genotypes of mice treated with drug or vehicle as shown and processed at 6 weeks of age. Scale bar: 200 µm

f-h) Quantitative PCR analysis for skeletal muscle-specific genes, *Atp2a1*, *Mylpf*, and *Tnni2* in hearts of the genotypes shown treated with drug or vehicle and analyzed at 6 weeks of age. Gene expression was normalized using 18S ribosomal RNA. n=2-4 mice per group. *$p<0.05$.
Data are presented as mean +/- SEM. Two-way ANOVA was used for statistical analysis.

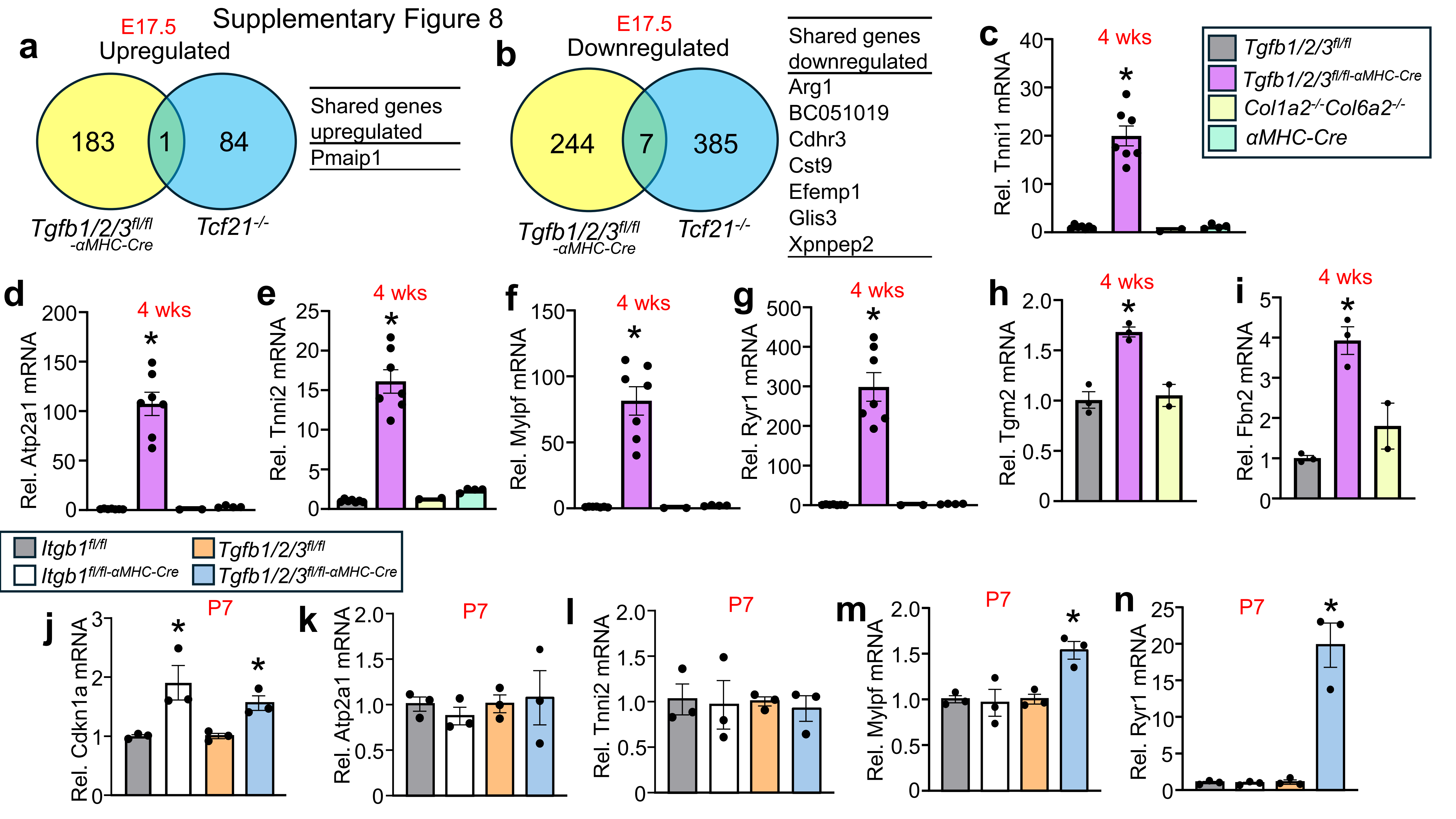
Supplementary Figure 8
a
E17.5
Upregulated
183
1
84
Tgfb1/2/3fl/fl -αMHC-Cre
Tcf21-/-
Shared genes upregulated
Pmaip1
b
E17.5
Downregulated
244
7
385
Tgfb1/2/3fl/fl -αMHC-Cre
Tcf21-/-
Shared genes downregulated
Arg1
BC051019
Cdhr3
Cst9
Efemp1
Glis3
Xpnpep2
c
4 wks
Rel. Tnni1 mRNA
Tgfb1/2/3fl/fl
Tgfb1/2/3fl/fl-αMHC-Cre
Col1a2-/-Col6a2-/-
αMHC-Cre
d
4 wks
Rel. Atp2a1 mRNA
e
4 wks
Rel. Tnni2 mRNA
f
4 wks
Rel. Mylpf mRNA
g
4 wks
Rel. Ryr1 mRNA
h
4 wks
Rel. Tgm2 mRNA
i
4 wks
Rel. Fbn2 mRNA
Itgb1fl/fl
Itgb1fl/fl-αMHC-Cre
Tgfb1/2/3fl/fl
Tgfb1/2/3fl/fl-αMHC-Cre
j
P7
Rel. Cdkn1a mRNA
k
P7
Rel. Atp2a1 mRNA
l
P7
Rel. Tnni2 mRNA
m
P7
Rel. Mylpf mRNA
n
P7
Rel. Ryr1 mRNA

**Supplementary Fig. 8. Cardiomyocyte-specific loss of *Tgfb1/2/3* genes reprograms the fibroblast and leads to aberrant cardiomyocyte differentiation**

a) Venn diagram comparison of differentially upregulated genes from hearts of *Tgfb1/2/3*$^{fl/fl\text{-}\alpha MHC\text{-}Cre}$ mice (n=3) in comparison to differentially upregulated genes from hearts of*Tcf21*$^{-/-}$ mice taken at E17.5 of gestation. RNA was extracted from flash frozen hearts from *Tcf21*$^{-/-}$ mice (n=2) and hearts from control, *Tcf21*$^{+/+}$ littermates (n=3) at E17.5. Differentially regulated genes were determined by comparison of gene expression of both *Tcf21*$^{-/-}$ and *Tgfb1/2/3*$^{fl/fl\text{-}\alpha MHC\text{-}Cre}$ to their respective control littermates and overlapping differentially regulated genes were compared. The 2 genotypes shared only 1 overlapping differentially upregulated gene.

b) Venn diagram comparison of differentially downregulated genes from hearts of *Tgfb1/2/3*$^{fl/fl\text{-}\alpha MHC\text{-}Cre}$ mice (N=3) in comparison to differentially downregulated genes from hearts of*Tcf21*$^{-/-}$ mice taken at E17.5 of gestation. RNA was extracted from flash frozen hearts from *Tcf21*$^{-/-}$ mice (n=2) and hearts from control, *Tcf21*$^{+/+}$ littermates (n=3) at E17.5. The 2 genotypes shared only 7 overlapping differentially downregulated genes.

c-i) Quantitative PCR analysis of *Tnni1*, *Atp2a1*, *Tnni2*, *Mylpf*, *Ryr1*, *Tgfm2*, and *Fbn2* from hearts of *Tgfb1/2/3*$^{fl/fl}$, *Tgfb1/2/3*$^{fl/fl\text{-}\alpha MHC\text{-}Cre}$, *Col1a2*$^{-/-}$*Col6a2*$^{-/-}$, and αMHC-Cre mice at 4 weeks of age. Expression was normalized to 18S ribosomal RNA expression. n=2-7 mice per group. *$p<0.05$.

j-n) Quantitative PCR analysis of *Cdkn1a*, *Atp2a1*, *Tnni2*, *Mylpf*, and *Ryr1* from hearts of *Itgb1*$^{fl/fl}$, *Itgb*$^{fl/fl\text{-}\alpha MHC\text{-}Cre}$, *Tgfb1/2/3*$^{fl/fl}$, and *Tgfb1/2/3*$^{fl/fl\text{-}\alpha MHC\text{-}Cre}$ mice at postnatal day 7 (P7). Expression was normalized to 18S ribosomal RNA expression and to their respective control littermate. n=3 mice per group. *$p<0.05$.
c-n. Data presented as mean +/- SEM. One-way ANOVAs used for statistical analysis.

Supplementary Figure 9

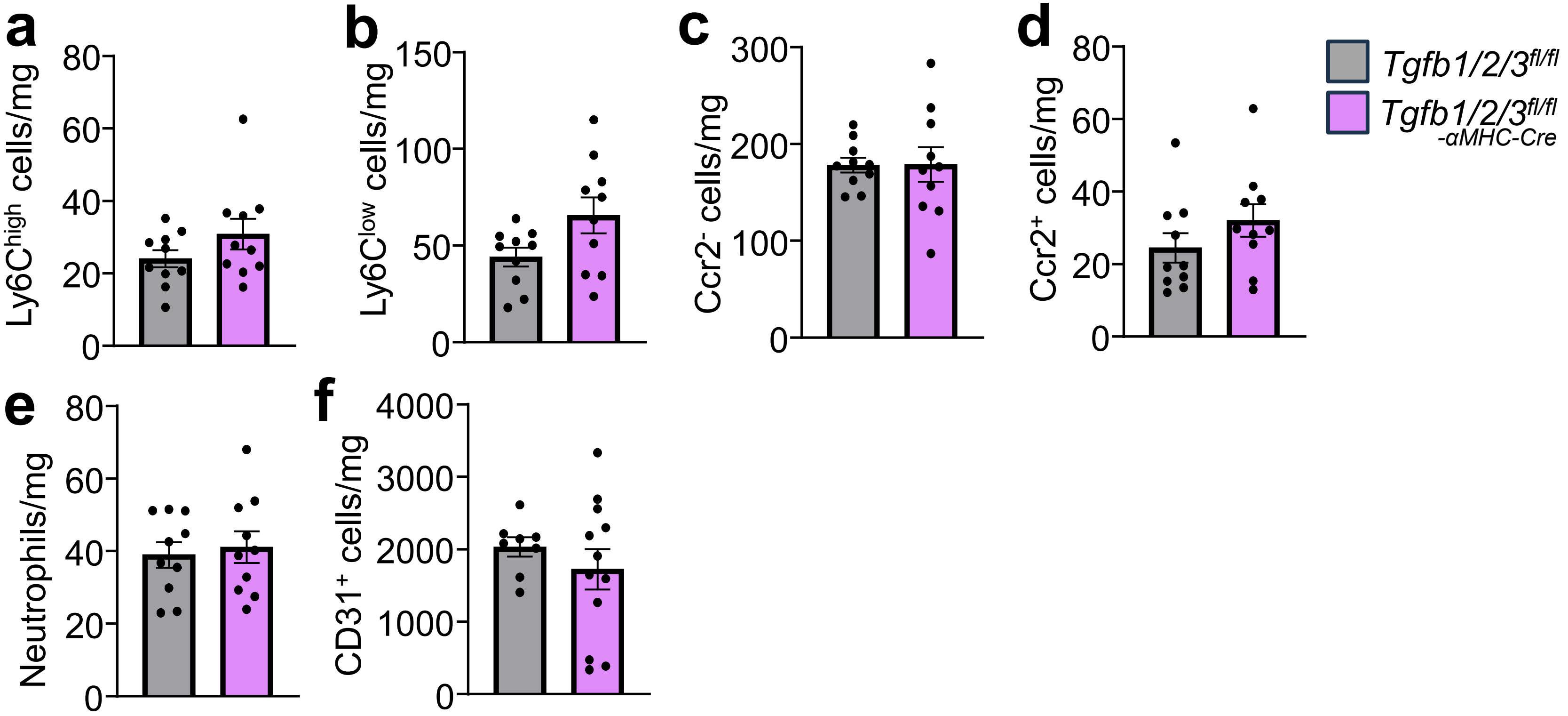


**Supplementary Fig. 9. Analysis of inflammatory cell response in P7 hearts from *Tgfb1/2/3$^{fl/fl-αMHC-Cre}$* and *Tgfb1/2/3$^{fl/fl}$* mice.**

a) Flow cytometric quantification of $CD11b^+F4/80^+Ly6C^{high}$ monocytes ($Ly6C^{high}$) from P7 hearts of *Tgfb1/2/3$^{fl/fl-αMHC-Cre}$* mice and *Tgfb1/2/3$^{fl/fl}$* littermates controls. All cell counts were normalized to tissue mass. n=10 per group.

b) Flow cytometric quantification of $CD11b^+F4/80^+Ly6C^{low}$ monocytes ($Ly6C^{low}$) from P7 hearts of *Tgfb1/2/3$^{fl/fl-αMHC-Cre}$* mice and *Tgfb1/2/3$^{fl/fl}$* littermates. n=10 per group.

c) Flow cytometric quantification of $Ccr2^-$ macrophages from P7 hearts of *Tgfb1/2/3$^{fl/fl-αMHC-Cre}$* mice and *Tgfb1/2/3$^{fl/fl}$* littermates. n=10 per group.

d) Flow cytometric quantification of $Ccr2^+$ macrophages from P7 hearts of *Tgfb1/2/3$^{fl/fl-αMHC-Cre}$* mice and *Tgfb1/2/3$^{fl/fl}$* littermates. n=10 per group.

e) Flow cytometric quantification of neutrophils from P7 hearts of *Tgfb1/2/3$^{fl/fl-αMHC-Cre}$* mice and *Tgfb1/2/3$^{fl/fl}$* littermates. n=10 per group.

f) Flow cytometric quantification of CD31+ cells from P7 hearts of *Tgfb1/2/3$^{fl/fl-αMHC-Cre}$* mice and *Tgfb1/2/3$^{fl/fl}$* littermates. n=10 per group.

Data are presented as mean +/- SEM. Student's T-test was used for statistical analysis.

**Supplementary Table 1. List of significantly increased and decreased genes in isolated fibroblasts from adult hearts deleted for *Tgfb1/2/3* with the αMHC-Cre transgene.**

| Gene ID | Gene name | FC | Gene function |
|---|---|---|---|
| Crlf1 | cytokine receptor-like factor 1 | 9.40 | Fibroblast-induced in response to cytokine signaling, promotes cardiac fibrosis via ERK1/2 signaling pathway |
| Ddah1 | dimethylarginine dimethylaminohydrolase 1 | 8.03 | Breaks down asymmetric dimehtylarginine (ADMA) that inhibits nitric oxide production. DDAH inhibition reduces fibroblast induced collagen production |
| Notch4 | notch 4 | 6.19 | Promotes myofibroblast differentiation and collagen synthesis potentially independent of TGFβ |
| Wisp2 | WNT1 inducible signaling pathway protein 2 | 5.40 | Highly expressed in mesenchymal stem cells. Secreted WISP-2 drives proliferation of mesenchymal precursor cells and maintains them in an undifferentiated state |
| Bdnf | brain derived neurotrophic factor | 5.04 | Expressed by fibroblasts, especially cancer associated fibroblasts. Plays role in cell growth, survival, and differentiation. |
| Angptl7 | angiopoietin-like 7 | 5.00 | Extracellular protein that is involved in ECM organization, especially in the eye. Overexpression results in reduced expression of collagens and fibronectin as well as altered fibronectin assembly and secretion. |
| Cx3cl1 | chemokine (C-X3-C motif) ligand 1 | 4.52 | Fractalkine; expressed by fibroblasts during wound healing or fibrosis to attract immune cells including macrophages |
| Gdf6 | growth differentiation factor 6 | 4.29 | Regulates function and differentiation of mesenchymal stem cells |
| Ereg | epiregulin | 3.96 | Upregulated in cancer associated fibroblasts (CAFs) where it drives tumor cell proliferation, migration, and invasion within the tumor microenvironment |
| Cst6 | cystatin E/M | 3.37 | Associated with cancer-associated fibroblasts as downregulation results in reduced tumor progression and metastasis. Functions as cysteine protease inhibitor that binds to and inactivates cysteine cathepsins which are involved in degradation of ECM proteins including collagen, elastin, and proteoglycans. Imbalance between cathepsins and their inhibitors can result in dysregulation of ECM remodeling. |
| Ahsg | alpha-2-HS-glycoprotein | 3.03 | Glycoprotein generally produced by the liver. Associated with coronary atherosclerosis. Hypothesized to have role in tissue development. Upregulated in isolated cardiac fibroblasts from hearts of *Tgfbr1/2-Tcf21MCM/+* mice [3]. |
| Fetub | fetuin beta | 2.68 | Protease inhibitor. Upregulated in isolated cardiac fibroblasts from hearts of Tgfbr1/2-Tcf21MCM/+ mice [3]. |
| Nbl1 | neuroblastoma, suppression of tumorigenicity 1 | 2.60 | Secreted protein that binds to and antagonizes BMP. Thus, maybe important during development. Reduces scar formation in corneal fibrosis. |
| Cst3 | cystatin C | 2.46 | Secreted protein that has inhibitory effect on fibroblasts by suppressing their proliferation and activation especially in the context of fibrosis |
| Ccrl1 | chemokine (C-C motif) receptor-like 1 | 2.45 | Atypical chemokine receptor that acts as a chemokine scavenger. Can regulate immune cell trafficking and organization |
| Meox1 | mesenchyme homeobox 1 | 2.28 | Promotes fibroblast activation and ECM production. High Meox1 expression associated with fibroblast activation |
| Sparc | secreted acidic cysteine rich glycoprotein | 2.09 | Promotes fibroblast proliferation, migration, activation, and ECM production |

| | | | |
|---|---|---|---|
| Bgn | biglycan | 2.03 | Small leucine rich proteoglycan involved in ECM organization and assembly. Biglycan associated with poor prognosis in certain cancers by transforming mesothelial cells to cancer associated fibroblasts which in turn promoted tumor proliferation, migration, and invasion. Exhibits immunosuppressive functions in certain cancers |
| Runx1 | runt related transcription factor 1 | 1.98 | Regulates myofibroblast differentiation; Runx1 expression associated with fibroblast proliferation and ECM production |
| Wipf3 | WAS/WASL interacting protein family, member 3 | 1.98 | Expressed by fibroblasts. Predicted to enable alpha tubulin and gamma tubulin binding activity. Modulates cytoskeletal dynamics in podocytes. |
| Id1 | inhibitor of DNA binding 1 | 1.96 | Negative regulator of fibrosis; suppresses excessive collagen production by fibroblasts in response to injury or TGFβ |
| Hes1 | hairy and enhancer of split 1 (Drosophila) | 1.90 | Maintains fibroblast quiescence by acting as a transcriptional repressor to prevent premature senescence and differentiation. |
| Sfrp1 | secreted frizzled-related protein 1 | 1.83 | Inhibits fibroblast proliferation, migration and activation in many contexts including wound healing and cancer; interacts with Wnt signaling |
| Tmem254c | transmembrane protein 254c | -16.5 | Expressed in fibroblasts. Associated with arrhythmogenic cardiomyopathy. |
| Cxcl13 | chemokine (C-X-C motif) ligand 13 | -10.9 | Produced by fibroblasts and functions as a B cell chemoattractant. Fibroblasts expressing CXCL13 contribute to immune cell recruitment and inflammation. |
| Lag3 | lymphocyte-activation gene 3 | -9.96 | Cell surface protein. Expression of Lag3 on fibroblasts may regulate their function through influencing their activation and extracellular matrix production |
| Pttg1 | pituitary tumor-transforming gene 1 | -9.81 | Pituitary tumor-transforming gene 1. Overexpression induces chromatin instability resulting in cell senescence and growth arrest |
| Trpc3 | transient receptor potential cation channel, subfamily C, member 3 | -8.82 | Regulates cardiac fibroblast proliferation and differentiation through calcium influx |
| Efemp1 | epidermal growth factor-containing fibulin-like extracellular matrix protein 1 | -7.36 | Encodes fibulin-3, an extracellular matrix glycoprotein expressed by fibroblasts, particularly in senescent or quiescent fibroblasts. Regulates ECM structure and function |
| Ccl11 | chemokine (C-C motif) ligand 11/Eotaxin | -7.04 | Profibrogenic cytokine that selectively increases fibroblast proliferation, matrix metalloprotease 2 activity, and collagen synthesis |
| Mme | membrane metallo endopeptidase (Neprilysin) | -6.77 | MME-positive fibroblasts constitute a subpopulation of cancer-associated fibroblasts that promote growth and invasion. Degrades bioactive peptides and growth factors in the ECM. |
| Bcam | basal cell adhesion molecule | -6.52 | Membrane protein expressed by fibroblasts and regulates cell adhesion to the extracellular matrix by interacting with laminins including Lama5. |
| Htra1 | HtrA serine peptidase 1 | -5.13 | Serin protease. Expression induced by TGFβ1. *In vitro* suppression of HTRA1 inhibits conversion of cardiac fibroblasts into myofibroblasts. HTRA1 knockdown results in Collagen I retention in the ER resulting in ER stress and lysosomal degradation of collagen |
| Sorbs2 | sorbin and SH3 domain containing 2 | -4.64 | Can contribute to premature senescence in primary human fibroblasts. |
| Igfbp3 | insulin-like growth factor binding protein 3 | -4.11 | Promotes maturation of ex vivo embryonic murine cardiomyocytes and in vitro hiPSC cardiomyocytes. Plays critical role in myofibroblast differentiation in diseased tissue |

| | | | |
|---|---|---|---|
| Asb2 | ankyrin repeat and SOCS box-containing 2 | -4.08 | E3 ubiquitin ligase specificity subunit. Regulates hematopoietic differentiation and myogenic differentiation through targeted degradation of filamins that alter cell adhesion, spreading, and actin remodeling. |
| Meis3 | Meis homeobox 3 | -3.95 | Homeobox protein and transcriptional regulator. Involved in regulating developmental differentiation and promotes cell survival |
| Gpc3 | glypican 3 | -3.83 | Cell-surface proteoglycan. Upregulated in cancer fibroblasts promoting cellular growth and migration. Binds Hedgehog (Hh), Wnt, and fibroblast growth factor 2 via its heparin sulphate glycan chains |
| Tmeff2 | transmembrane protein with EGF-like and two follistatin-like domains 2 | -3.46 | Overexpression inhibits fibroblast proliferation which could thereby function to limit tumor prognosis |
| Amigo2 | adhesion molecule with Ig like domain 2 | -3.30 | Induced by TGFβ. Upregulated in cancer associated fibroblasts and induces release of tumor-promoting secretomes. Mediates cancer cell-endothelial cell adhesion to drive metastases |
| Olfml2a | olfactomedin-like 2A | -3.13 | Secreted glycoprotein expressed in fibroblasts. Upregulated in cancers to promote proliferation, associated with poor prognosis |
| S1pr3 | sphingosine-1-phosphate receptor 3 | -3.10 | Highly expressed in fibroblasts. Promotes fibrosis. |
| Il18 | interleukin 18 | -2.86 | Pro-inflammatory cytokine produced by mesenchymal cells |

Selected genes dysregulated in fibroblasts of *Tgfb1/2/3fl/fl-αMHC-Cre* mice at 6 weeks of age that were not dysregulated in single nuclear sequencing analysis of fibroblasts in pathological cardiomyopathies [66, 67]. Descriptions of the "Gene Function" and association to fibroblast biology were summaries generated by ChatGPT that we verified with independent searches. FC, fold changes each had highly significant P values

**Supplementary Table 2.** Comparison of altered gene expression between hearts of *Tgfb1/2/3*$^{fl/fl\text{-}\alpha MHC\text{-}Cre}$ mice and 4 other models with deletion of chromatin regulators that show ectopic cardiac gene induction

| Gene | *Tgfb123*$^{fl/fl\text{-}}$ *αMHC-Cre* (6 wks) | *Jarid2*$^{fl/fl\text{-}}$ *αMHC-Cre* (7 mos) | *Hdac1/2*$^{fl/fl\text{-}}$ *αMHC-Cre* (P8) | *Chd4*$^{fl/fl\text{-}}$ *αMHC-Cre* (2 wks) | *Ezh2*$^{fl/fl\text{-}Mef2cAFH\text{-}}$ *Cre* (Adult) |
|---|---|---|---|---|---|
| *Atp2a1* | 18.0 | 2.00 (P10) | | 48.12 | 22.2 |
| *Mybph1* | 2.30 | 8.10 | | | 76.2 |
| *Mylpf* | 14.0 | | 2.30 | 32.7 | 167 |
| *Tnni2* | 13.0 | | 111 | 12.9 | 31.7 |
| *Acta1* | 4.96 | | | 12.7 | 9.99 |
| *Tnnt3* | 4.92 | | | 78.3 | 254 |
| *Casq1* | 4.43 | | 2.50 | 15.5 | 1.82 |
| *Ctgf* | 3.73 | 5.06 | | 2.08 | 3.21 |
| *Ankrd1* | 2.50 | 2.50 | | 2.06 | 7.14 |
| *Myl4* | 2.00 | | 8.00 | | 18.9 |
| *Myl7* | 1.90 | | 97.0 | -2.23 | 25.7 |
| *Cdkn1a* | 18.3 | 2.75 | 2.10 | | 1.67 |
| *Efhd2* | 1.71 | 2.25 | 2.80 | 2.46 | 1.82 |
| *Nppa* | 6.80 | 13.2 | 4.00 | | 17.3 |
| *Nppb* | 4.70 | 5.50 | | 2.38 | 6.82 |
| *Sfrp1* | 2.70 | 2.70 | 8.00 | | 1.81 |
| *Sdc4* | 2.59 | 1.55 | 2.60 | | 1.89 |
| *Tpm2* | 2.01 | | 2.50 | 3.39 | 3.31 |
| *Thrsp* | 10.7 | | 2.10 | | 2.65 |
| *Kcnd2* | -2.90 | -3.80 | -2.27 | | -4.17 |
| *Tmem150c* | -2.10 | -2.10 | | | |
| *Klh133* | -2.50 | -2.50 | | | |
| *Kcnd2* | -2.90 | -3.80 | -2.27 | | -4.17 |
| *Tac1* | -1.70 | -3.60 | | | 5.23 |
| *Mme* | -2.00 | -3.70 | | | -2.13 |
| *Acot3* | -1.90 | -3.30 | -2.13 | | -5.00 |
| *Kcnv2* | -1.60 | -3.40 | | | -3.37 |
| *Clcn1* | -1.70 | -3.30 | | | -2.00 |
| *Fbp2* | -4.00 | -2.00 | | | -1.88 |
| *Gria3* | 4.10 | 2.30 | 4.90 | 2.02 | |
| *Spp1* | 1.70 | 4.30 | 5.70 | 41.9 | |
| *Gpx3* | 2.84 | 1.89 | 3.70 | | 2.29 |

The numbers show fold change in mRNA expression of each indicated gene versus wildtype controls, and negative numbers indicate reduced expression. Age of mice is given from hearts of each of the 5 cardiac-specific gene-deleted mice in the top column. *Tgfb1/2/3*$^{fl/fl\text{-}\alpha MHC\text{-}Cre}$ mice compared to hearts of *Jarid2*$^{fl/fl\text{-}\alpha MHC\text{-}Cre}$ [72], *Hdac1/2*$^{fl/fl\text{-}\alpha MHC\text{-}Cre}$ [71], *Chd4*$^{fl/fl\text{-}\alpha MHC\text{-}Cre}$ [73], and *Ezh2*$^{flfl\text{-}med2cAFH\text{-}Cre}$ [70], mice. The top portion of the listed genes represents skeletal muscle-specific gene expression observed in hearts of these mice. The bottom portion of the table (divided by the bold line) represents other overlapping genes including genes involved in cardiac development. Abbreviation: P, postnatal; mos, months; wks, weeks.

**Supplementary Table 3.** Downregulation of mitochondrial genes in hearts from *Tgfb1/2/3* deleted mice suggesting less differentiation

| Gene | P-value adj | Ave log2 FC downreg |
|---|---|---|
| *Atp5c1* | 0 | 0.83 |
| *Atp5f1* | 0 | 1.08 |
| *Ndufab1* | 0 | 2.01 |
| *Ndufa12* | 0 | 2.16 |
| *Ndufv2* | 0 | 1.13 |
| *Cox6a1* | 0 | 1.80 |
| *Hspd1* | 0 | 2.39 |
| *Higd1a* | 0 | 2.39 |
| *Ndufb5* | 0 | 2.58 |
| *Acadvl* | 4.37E-289 | 2.55 |
| *Cox8a* | 8.91E-289 | 2.42 |
| *Ech1* | 1.52E-276 | 2.12 |
| *Atp5g2* | 8.74E-257 | 1.77 |
| *Atp5h* | 3.23E-245 | 1.87 |
| *Ndufa8* | 2.73E-244 | 3.36 |
| *Cycs* | 1.14E-231 | 3.29 |
| *Ndufb3* | 2.30E-222 | 1.68 |
| *Ndufa7* | 3.02E-219 | 3.13 |
| *Mtch1* | 1.18E-216 | 1.40 |
| *Ndufb1-ps* | 2.93E-204 | 1.75 |
| *Ndufb2* | 6.63E-196 | 2.29 |
| *Cox6b1* | 6.32E-189 | 2.15 |
| *Ndufs2* | 6.52E-189 | 1.61 |
| *Cox5a* | 2.34E-186 | 1.26 |
| *Ndufc2* | 1.29E-183 | 3.89 |
| *Ndufv1* | 3.74E-178 | 1.46 |
| *Idh3g* | 5.86E-177 | 2.10 |
| *Acads* | 1.31E-175 | 3.04 |
| *Cox7c* | 1.10E-168 | 0.78 |
| *Ndufs7* | 6.62E-162 | 2.25 |
| *Tomm20* | 3.16E-148 | 1.90 |
| *Atp5j* | 1.57E-116 | 2.27 |
| *Bax* | 5.36E-76 | 2.82 |

List of mitochondrial genes used to generate a composite score of mitochondrial maturation from single nuclear RNA sequencing of hearts from *Tgfb1/2/3*$^{fl/fl\text{-}\alpha MHC\text{-}Cre}$ mice and *Tgfb1/2/3*$^{fl/fl}$ control mice at P7. n=2 per group. $p<0.05$, fold change (FC) <-1.5.

**Supplementary Table 4.** TGFβ TG heart gene expression changes at 6 months of age compared to control

| Symbol | Name | Fold change |
|---|---|---|
| *Mfap4* | microfibrillar-associated protein 4 | 4.248 |
| *Sfrp2* | secreted frizzled-related protein 2 | 2.893 |
| *Adamtsl2* | ADAMTS-like 2 | 2.155 |
| *Lox* | lysyl oxidase | 2.100 |
| *Itgbl1* | integrin, beta-like 1 | 2.022 |
| *Txnip* | thioredoxin interacting protein | 1.934 |
| *Pi16* | peptidase inhibitor 16 | 1.832 |
| *Angptl4* | angiopoietin-like 4 | 1.781 |
| *Col8a1* | Collagen, type VIII, alpha 1 | 1.724 |
| *Nr4a1* | nuclear receptor subfamily 4, group A, member 1 | 1.672 |
| *Pmepa1* | prostate transmembrane protein, androgen induced 1 | 1.640 |
| *Myc* | myelocytomatosis oncogene | 1.599 |
| *Serpina3n* | serine (or cysteine) peptidase inhibitor, clade A, member 3N | 1.548 |
| *Ctgf* | connective tissue growth factor | 1.537 |
| *Angptl7* | angiopoietin-like 7 | 1.519 |
| *Bgn* | biglycan | 1.502 |
| *Neb* | Nebulin | 1.498 |
| *Nrg1* | neuregulin 1 | 1.383 |
| *Ncam1* | neural cell adhesion molecule 1 | 1.375 |
| *Mrc1* | mannose receptor, C type 1 | -1.335 |
| *Cd163* | CD163 antigen | -1.358 |
| *Fbxl21* | F-box and leucine-rich repeat protein 21 | -1.423 |
| *Cyp2e1* | cytochrome P450, family 2, subfamily e, polypeptide 1 | -1.502 |
| *Mstn* | myostatin | -1.514 |
| *Cdkn1a* | cyclin-dependent kinase inhibitor 1A (P21) | -1.527 |
| *Acta1* | actin, alpha 1, skeletal muscle | -1.536 |
| *Clec10a* | C-type lectin domain family 10, member A | -1.630 |
| *Efemp1* | epidermal growth factor-containing fibulin-like extracellular matrix protein 1 | -1.660 |
| *Wtap* | Wilms' tumour 1-associating protein | -1.702 |
| *Myl4* | myosin, light polypeptide 4 | -1.751 |
| *Ccl11* | chemokine (C-C motif) ligand 11 | -2.143 |

All fold change values were $<0.05$ P value.  Negative values reflect reduced expression.
